# Contemporary HIV-1 consensus Env with redesigned hypervariable loops promote antibody binding

**DOI:** 10.1101/2023.11.19.567729

**Authors:** Hongjun Bai, Eric Lewitus, Yifan Li, Vincent Dussupt, Bonnie Slike, Letzibeth Mendez-Rivera, Annika Schmid, Lindsay Wieczorek, Victoria Polonis, Shelly J. Krebs, Julie A. Ake, Sandhya Vasan, M. Gordon Joyce, Samantha Townsley, Morgane Rolland

## Abstract

An effective HIV-1 vaccine must elicit broadly neutralizing antibodies (bnAbs) against the highly diverse Envelope glycoproteins (Env) present globally. Since Env with the longest hypervariable (HV) loops were more resistant to the cognate bnAbs than Env with shorter HV loops, we redesigned hypervariable loops for updated HIV-1 Env consensus sequences of subtypes B and C and circulating recombinant form AE (CRF01_AE). We reduced the length of V1HV, V2H, and V5HV while maintaining the integrity of the Env structure and glycan shield, and we modified V4HV to account for its diverse structural context. Redesiged HV loops consisted mainly of glycine and serine to limit strain-specific targeting. Redesigned consensus Env of subtype B or CRF01_AE demonstrated increased magnitude of binding responses to pooled plasma samples and representative bnAbs. Together with other antigen optimization techniques, consensus Env with redesigned hypervariable loops can improve future HIV-1 vaccine antigens to elicit bnAbs.

## Introduction

HIV-1 remains a global health priority. Vaccines have historically been proven to be the most powerful tool to fight against viruses. Despite many setbacks in HIV-1 vaccine efficacy trials, the search for a safe and effective HIV-1 vaccine should continue. Several unique HIV-1 features have hampered the development of an HIV-1 vaccine. The protein that allows HIV-1 entry, Env, is metastable, heavily glycosylated and highly diverse. To counteract HIV-1 diversity, broadly neutralizing antibodies (bnAbs) must be elicited by a vaccine to cross-react with a variety of Env^1,2^. The vaccine candidates tested in the nine HIV-1 vaccine efficacy trials conducted to date did not yield bnAbs. Structural biology studies have revealed the atomic details of the epitopes recognized by numerous HIV-1 bnAbs (e.g. as reviewed^3–5)^, providing a roadmap for designing antigens that could integrate these bnAb epitopes. Known HIV-1 bnAbs cover the Env surface^6^ and are grouped in categories based on the location of their epitopes, including Env variable loop 2 (V2)-apex antibodies, glycan supersite antibodies and CD4 binding site (CD4bs) antibodies^3–5^. Those bnAbs epitope are under functional constraints. For example, the V2-apex epitope shields the V3 loop and stabilizes the prefusion Env trimer^7,8^ and the CD4bs epitope is where the host receptor binds to^9^.

HIV-1 Env present five variable loops and four of them (V1, V2, V4, and V5) contain a hypervariable segment, called the hypervariable (HV) loop. These HV loops can be adjacent to epitope sites recognized by bnAbs. The V1 HV loop (V1HV) is near the glycan supersite epitope (target of 10-1074^10^, PGT135^11,12^, PGT128^11,13^ and 2G12^14^), the V2 HV loop (V2HV) is near the V2-apex epitope (target of PGT145^15–17^, VRC26.25^18^, PG9^19^ and PGDM1400^15^), and the V5 HV loop (V5HV) is near the CD4bs epitope (target of VRC01^20^ and 3BNC117^21^). These HV loops can occlude access to the adjacent bnAbs epitopes and can be targeted by antibodies to elicit strain-specific responses rather than the more functional bnAb specificities they are shielding. Antibodies targeting these three epitopes accounted for more than half of the neutralizing antibodies in two natural infection cohorts: they corresponded to 57% of dominant specificities of the top 42 neutralizers in the IAVI Protocol C^22^ and to 65% (33/51) of delineated epitope specificities in the RV217 cohort^23^. Thus, modifying hypervariable loops to optimize access to adjacent bnAb epitopes on Env could improve HIV-1 Env vaccine candidates.

Previous studies analyzed HIV-1 antigens in which either V1, V2 or both loops were removed^24–26^. These variable loop deletions refocused immune responses, yet they also revealed that the removal of entire loops can lead to protein misfolding. Other studies used natural sequences with short HV loops, especially shorter V1HV, as candidate antigens. Zolla-Pazner and collaborators tested V1V2 from strain ZM53, which had a short V1, on scaffold proteins as vaccine antigens^27,28^. We selected a CRF01_AE strain with short V1HV and V2HV to be loaded on a self-assembling nanoparticle, V1V2-SHB-SAPN^29^. However, a natural sequence does not necessarily have ideal bnAb epitopes as well as ideal HV loops for eliciting bnAbs.

To overcome the limitations of natural HIV-1 sequences, we propose to systematically redesign HV loops to optimize access to bnAb epitopes. Our goals were to reduce the shielding of bnAb epitopes caused by HV loops and to limit the targeting of HV loops by strain specific antibodies. We used AlphaFold2^30^, a tool that predicts protein structure and, thereby, allows systematically redesign HV loops. We analyzed publicly available circulating subtype B and C and Circulating Recombinant Form (CRF) 01_AE sequences to infer naturally occurring loop lengths and AlphaFold2^30^ to predict protein structure to maintain structural integrity, loop anchoring, and the glycan shield. We redesigned Env HV loops for updated consensuses of subtypes B and C and CRF01_AE (Lewitus et al., under review) that we recently created using publicly available^31^ independent sequences sampled only since 2010 and tested bnAb accessibility of our redesigned sequences against unmodified Env consensus proteins. We used consensus sequences because they are a better representation of HIV-1 diversity than any natural sequence and are thus optimal as vaccine candidates^32–35^. We hypothesize that subtype consensus sequences with redesigned HV loops will reduce the shielding of bnAb epitopes caused by HV loops and limit the targeting of HV loops by strain specific antibodies.

## Results

### Short HV loops were rare for subtypes B, C and CRF01_AE

We curated publicly available circulating HIV-1 sequences to obtain a dataset of 4847 independent Env sequences sampled from people live with HIV (PLWH) between 1979 and 2020 (Lewitus et al., under review). We focused on subtype B (n=2495), subtype C (n=1503) and CRF01_AE (n=849) (Table S1) sequences since these subtypes/CRF are most frequently sequenced, thereby enabling robust analyses. We analyzed the distribution of HV loop lengths and the number of potential N-linked glycosylation sites (PNGS) in four HV loops (Fig. 1, Tables S1 and S2). We focused on the HV loops, which correspond to a portion of the variable loops: V1HV, sites 132-152 within sites 131-157 (V1); V2HV, sites 185-190 within sites 157-196 (V2); V4HV, sites 395-412 within sites 385-418 (V4); V5HV, sites 460-467 within sites 460-470 (V5). V3 was omitted because it does not contain an HV segment. The location of the HV loops, as well as adjacent bnAb epitopes, are shown on the structure in Fig. 1A.

**Fig 1.**
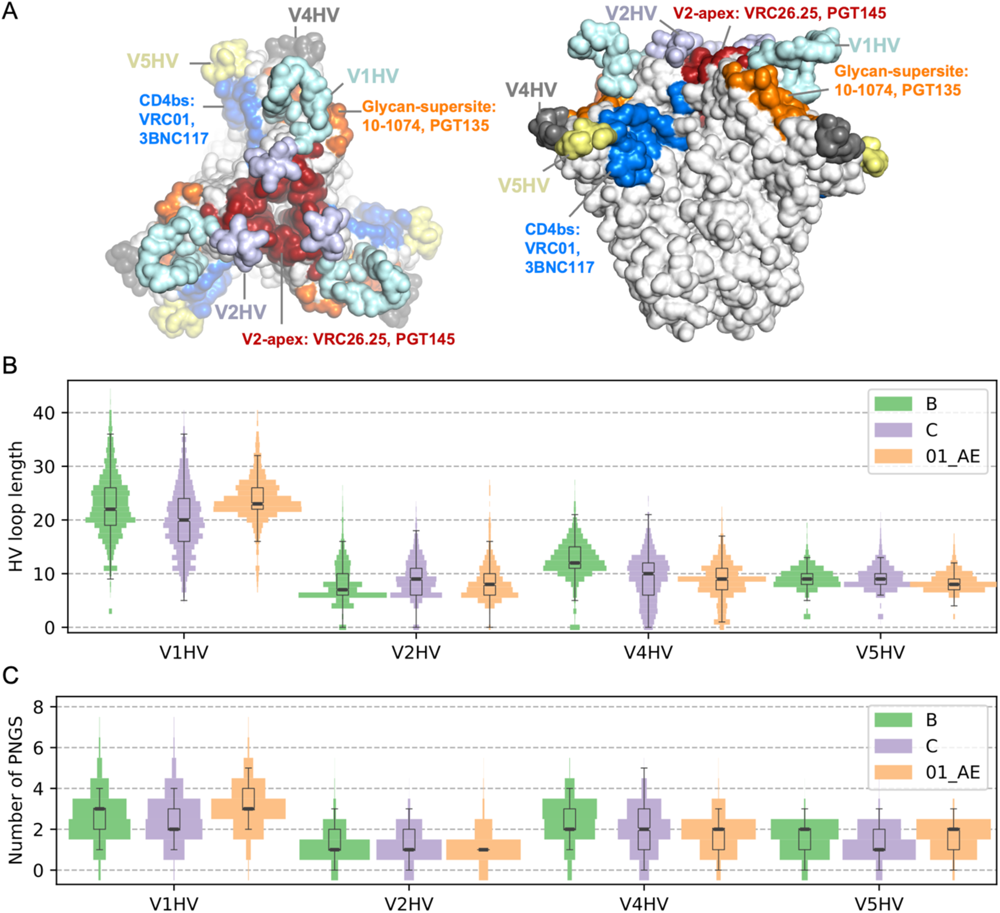
Characteristics of hypervariable (HV) loops. (A) Location of HV loops and related HIV-1 bnAb epitopes on Env prefusion structure. For subtypes B and C and CRF01_AE Env sequences, (B) the distribution of HV loop length for V1, V2, V4 and V5 HV loops and (C) the distribution of the number of potential N-linked glycosylation sites (PNGS) are shown. The width at each number corresponds to the proportion of sequences in the subtype/CRF sample of sequences and is normalized for each HV loop. Key characteristics of the distribution, including median and 25th/75th percentiles, are depicted by box plots.

V1HV loops showed a median length of 20 to 23 amino acids (AA), which was about twice as long as V2HV (7-9 AA), V4HV (9-12 AA), and V5HV (8-9 AA) loops (Fig. 1B, Table S1). The length distributions across subtypes/CRF were similar for V2HV and V5HV, while there were differences for V1HV and V4HV. CRF01_AE showed fewer short V1HV loops than subtype B or subtype C: there were 8% of CRF01_AE Env with V1HV ≤ 20 AA compared to 26% and 49% for B and C Env, respectively. Subtype B had fewer short V4HV loops than subtype C and CRF01_AE: there were 13% of subtype B Env with V4HV ≤ 10 AA compared to 49% and 65% for C and CRF01_AE Env, respectively. Similar patterns were observed for the number of PNGS on HV loops across subtypes/CRF (Fig. 1C, Table S2). Sequences with V1, V4, and V5 HV loops that were shorter than or equal to the 10th percentile were very rare representing 0.73%, 0.53%, and 0.32% of the dataset for subtypes B, C, and CRF01_AE, respectively. We also determined the loop lengths of sequences that have been used as inserts in previous vaccine efficacy trials or that are under consideration for future trials. We found that these insert sequences usually had one or more long HV-loops (Table 1)^36–45^. For example, the subtype C TV1 sequence which was included in the vaccine tested in the HVTN702^44^ vaccine efficacy trial had a V1HV loop that was 33 AA long while the median V1HV for subtype C is 20 AA. The insert Mosaic2.Env^39^ used in the Mosaico and Imbokodo vaccine efficacy trials had a V2HV that was twice as long (16 AA) as the median for subtypes B and C (7 and 9 AA, respectively). We identified some exceptions with short HV loops: the subtype A1 92RW020^40^ insert that was part of the trivalent vaccine tested in the HVTN505 trial and the subtype C CH505^43^ insert that has been undergoing early clinical testing.

**Table 1.**
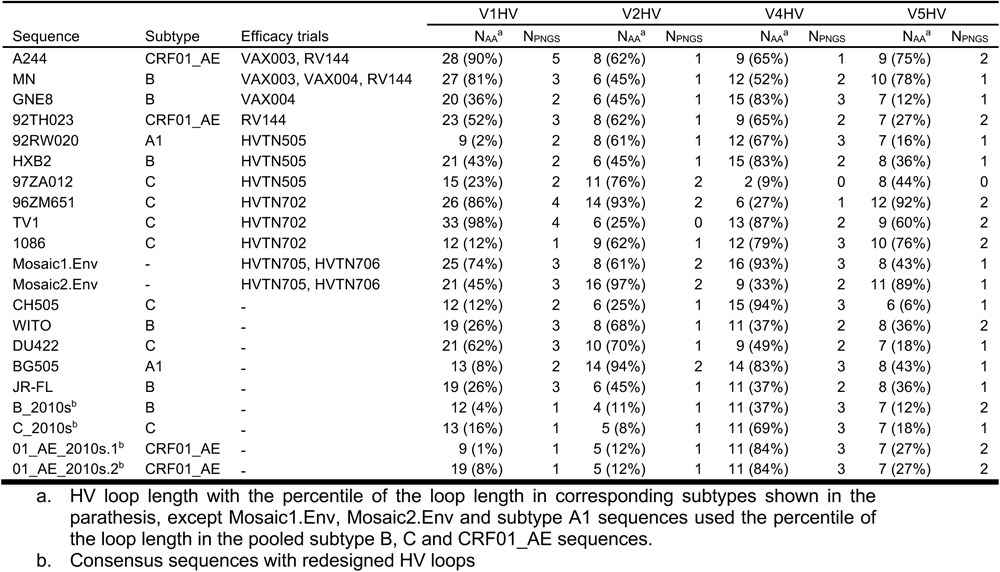
Comparison of redesigned HV loops to those found in vaccine candidates.

### Long HV loops associated with decreased neutralization sensitivity to bnAbs

The neutralization sensitivity to bnAbs was compared for HIV-1 Env with the shortest (≤ 5 percentile) and longest (≥ 95 percentile) HV loops. All neutralization and corresponding sequence data were obtained from CATNAP^46^. Six bnAbs belonging to three representative epitope groups were compared: V2-apex, PGT145 and VRC26.25; glycan supersite, 10-1074 and PGT135; and CD4bs, VRC01 and 3BNC117 (Fig. 2). To evaluate the effect of loop length on bnAb sensitivity, the cognate HV loops corresponding to each bnAb were analyzed and a distal HV loop was used as a control for each comparison. The virus strains with the longest cognate HV loops had significantly higher half maximal inhibitory concentration (IC50) than the sequences with the shortest HV loops (p-value ≤ 0.015, one-tailed Mann–Whitney U test) for the six bnAbs (Fig. 2), suggesting that longer HV loops obstruct bnAb epitope availability. When HV loops distant from the bnAb epitope were compared, there was no significant difference in sensitivity for short or long HV loops for five of the six bnAbs tested (p-value ≥ 0.207). For VRC01, sequences with shorter HV loops for both V2 and V5 were significantly associated with increased sensitivity when compared to sequences with longer HV loops when we compared the 5% of sequences with the longest and shortest HV loops (Fig. 2E); if using 10% or more as the threshold to compare the extremes of the distributions, the difference remained significant for V5HV (p ≤ 0.005) but not for V2HV (p ≥ 0.412) (Fig. S2E). The strongest impact was seen for V1HV loops and the neutralization sensitivity to 10-1074. HIV-1 Envs with the shortest V1HV loops had a median IC50 greater than or equal to 5000 times lower than the strains with the longest V1HV loops (0.02 μg/ml vs. ≥ 100 μg/ml, one-tailed Mann-Whitney U test p-value 0.00003). For V5HV loops, which are distant from the 10-1074 epitope, there was no significant difference in sensitivity when short and long loops were compared (one-tailed Mann-Whitney U test p-value 0.207). Importantly, these data masked some subtype-specific differences, as differences between short and long HV loops were not manifest for subtype B except for 10-1074 (Fig. S1). Differences for subtype C were generally statistically significant, except for 10-1074 (p=0.129) and PGT135 (p=0.128). We repeated the analysis using 10-25% as the threshold to compare the extremes of the distributions (Fig. S2). Differences between short/long adjacent HV loops remained statistically significant (p<0.045) except for PGT145 (p>0.101) and VRC26.25 when the cutoff was set as 25% (p=0.079). Overall, these results indicated that a long HV loop can diminish the virus’s sensitivity to a bnAb.

**Fig 2.**
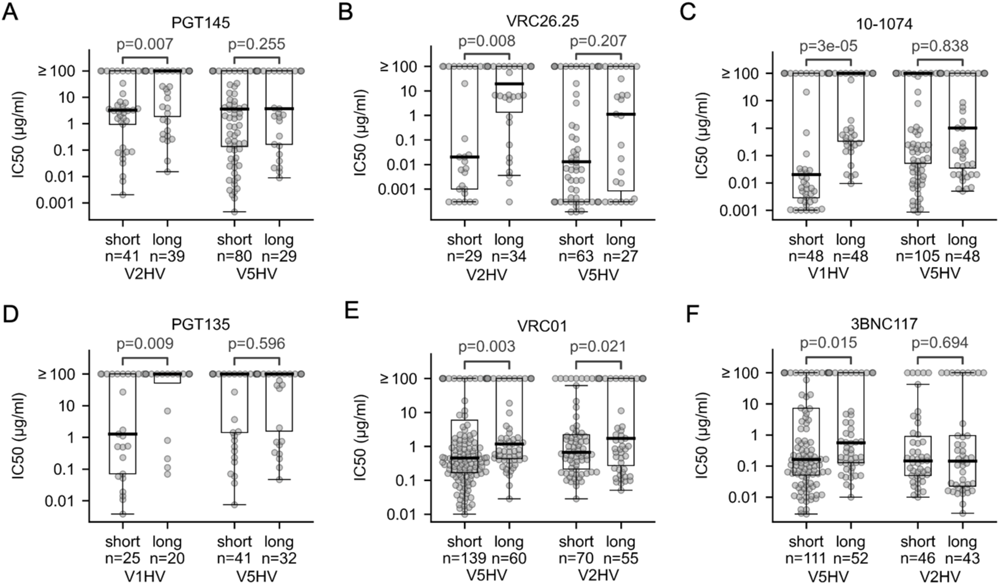
Comparison of the bnAb sensitivity for HIV-1 Env with short (≤ 5 percentile) or long (≥ 95 percentile) hypervariable (HV) loops. For each panel, comparisons of IC50 values for short and long HV loops are shown for the HV loop adjacent to (left) and remote from (right) the Ab epitope: PGT145 (A) and VRC26.25 (B), two V2 apex-targeting antibodies whose epitopes are close to V2HV loops but distant from V5HV loops; 10-1074 (C) and the PGT135 (D), two glycan supersite antibodies whose epitopes are close to V1HV loops but far from V5HV loops; VRC01 (E) and 3BNC117 (F), two CD4 binding site antibodies with epitopes close to the V5HV loops but far from V2HV loops. Key characteristics of the distribution, including median and 25th/75th percentiles, are depicted by box plots. The p-values from non-paired, non-parametric Mann-Whitney U test are provided.

### Strategy to redesign HV loops

We redesigned the HV loops on Env to improve accessibility to typical bnAb epitopes. We used Env consensus sequences which were derived only from sequences that were sampled after 2010, thus reflecting currently circulating viruses (Lewitus et al., under review). Our goals were to: (1) make the loop as short as possible while maintaining structural integrity; (2) preserve the loop anchor sites, which are the relatively conserved start and end of the HV loops that form favorable interactions with the non-HV part of Env; and (3) maintain the glycan shield by ensuring there was at least one PNGS on each redesigned loop. Using the V1HV loop of subtype B as an example, we analyzed the alignment of circulating sequences (Fig. 3A) and the AlphaFold2 predicted structure of the corresponding consensus sequence (Fig. 3B). We identified the relatively conserved TDL motif (sites 132-134) as N-anchor sites and the relatively conserved MEKG motif (sites 149-152) as C-anchor sites, with all sites in between considered spacer sites. In the TDL motif, D133 may form favorable charged interactions with K151 in the C-anchor sites or with K154; L134 may form hydrophobic interactions with I154, L175, I323 and I326. In the MEKG motif, M149 may form hydrophobic interactions with I326 and I154; E150 may form a favorable charged interaction with R419. These anchor sites were retained since they did not directly contact with glycan supersite antibodies. We tested different spacer lengths iteratively using AlphaFold2 predicted structures and shortened the 15 AA spacer to 5 AAs. This resulted in the V1HV loop length changing from 22 AA (50^th^ percentile in subtype B sequences) to 12 AA (4^th^ percentile in subtype B sequences) (Fig. 3BC). The V1HV, V2HV and V5HV loops of subtypes B, C and CRF01_AE were redesigned with the same strategy and are shown in Table 2 and Fig. S2-S10.

**Fig 3.**
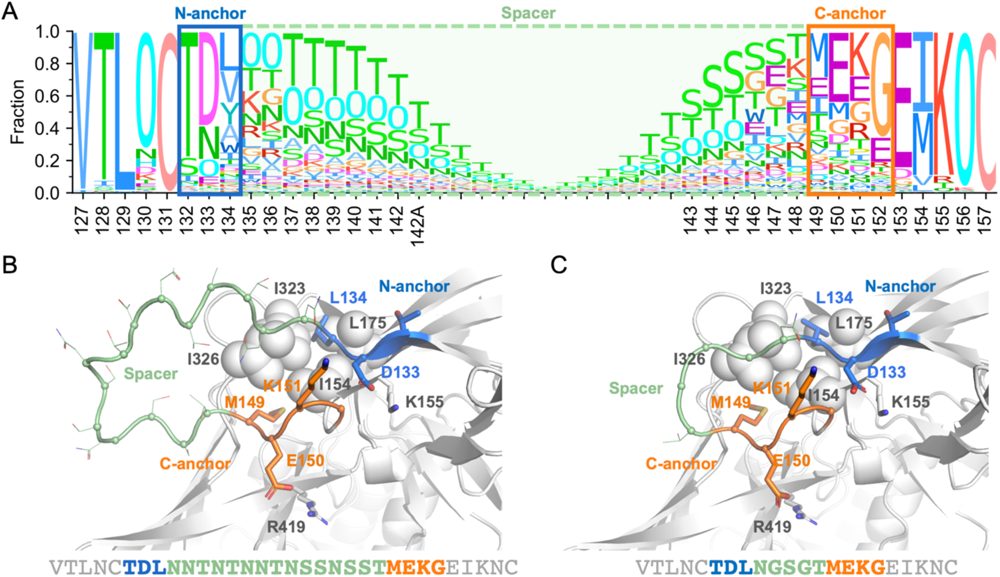
Redesigned V1 hypervariable loop of the subtype B Env consensus sequence. (A) Sequence logo of 2495 subtype B sequences around the V1HV loop, with the N-/C-anchors and spacer indicated by boxes. The height of each amino acid letter represents the frequency of that amino acid at a given position. The letter ‘O’ indicates a potential N-linked glycosylation site. The consensus B Env with unmodified V1HV loop (B) and redesigned V1HV loop (C) modeled by AlphaFold2 are shown with the N-anchor, C-anchor and spacer sites colored blue, orange and light green, respectively. Non-HV residues that interact with anchor sites are shown as spheres or sticks.

**Table 2.**
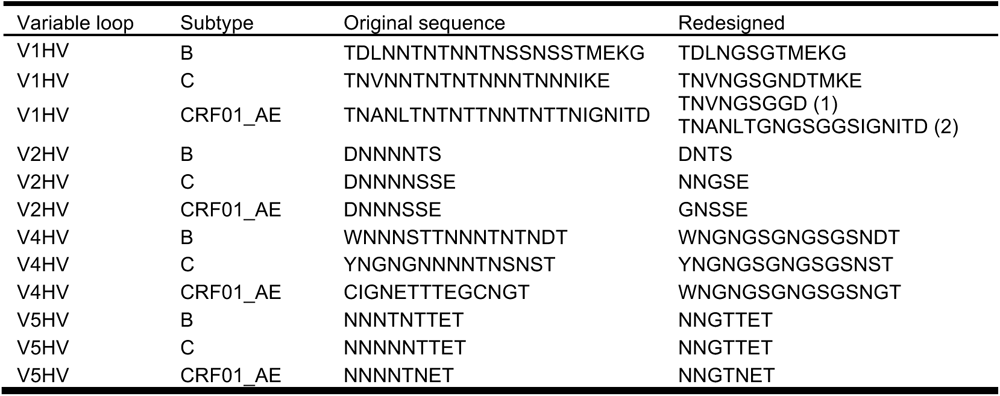
Sequences of redesigned HV loops.

Contrary to the strategy for V1HV, V2HV and V5HV, for V4HV loops we did not seek to obtain the shortest V4HV loops to accommodate its specific structural context. Sites adjacent to V4HV (non-HV sites within 10 Å of the HV loops, relative surface area > 0.30) were diverse, and no major bnAbs are known to target this segment of Env. For subtype C, the median Shannon entropy of V4HV adjacent sites was 2.4 bits compared to 0.8, 0.7 and 1.1 bits, respectively, for the sites adjacent to V1HV, V2HV and V5HV (Fig. 4A). This high Shannon entropy for V4HV loops indicated that there were 4.5 (2^2.4/1.1^) to 10.8 (2^2.4/0.7^) times more AA possibilities on average for surface sites adjacent to V4HV compared to the other HV loops. In addition, the distance encompassing spacer sites was larger for V4HV than for V1HV, V2HV and V5HVs (20.8 Å for V4HV versus 10.8, 7.9 and 9.4 Å for V1HV, V2HV and V5HV, respectively, based on the AlphaFold2 model of the subtype B consensus). This suggests that a short V4HV spacer may not be compatible with the structure. Hence, we chose a medium length for the V4HV loop; this loop length could also partially shield the highly diverse segment adjacent to the V4HV. For the subtype B consensus, we identified W395 as the N-terminal anchor site and ND (sites 411-412) as the C-terminal anchor sites. The spacer was replaced with an 11-AA linker (Fig. 4B-D). The redesigned linker contained two PNGS, which was the median number for V4HV of subtypes B and C, and CRF01_AE sequences (Fig. 1C). The V4HV loops of subtype C and CRF01_AE were redesigned with the same strategy (Fig. S11-S12). Across all redesigned HV loops, the spacer consisted mainly of glycine and serine which lack the side chains that are preferentially targeted by antibodies. All redesigned HV loops are listed in Table 2.

**Fig 4.**
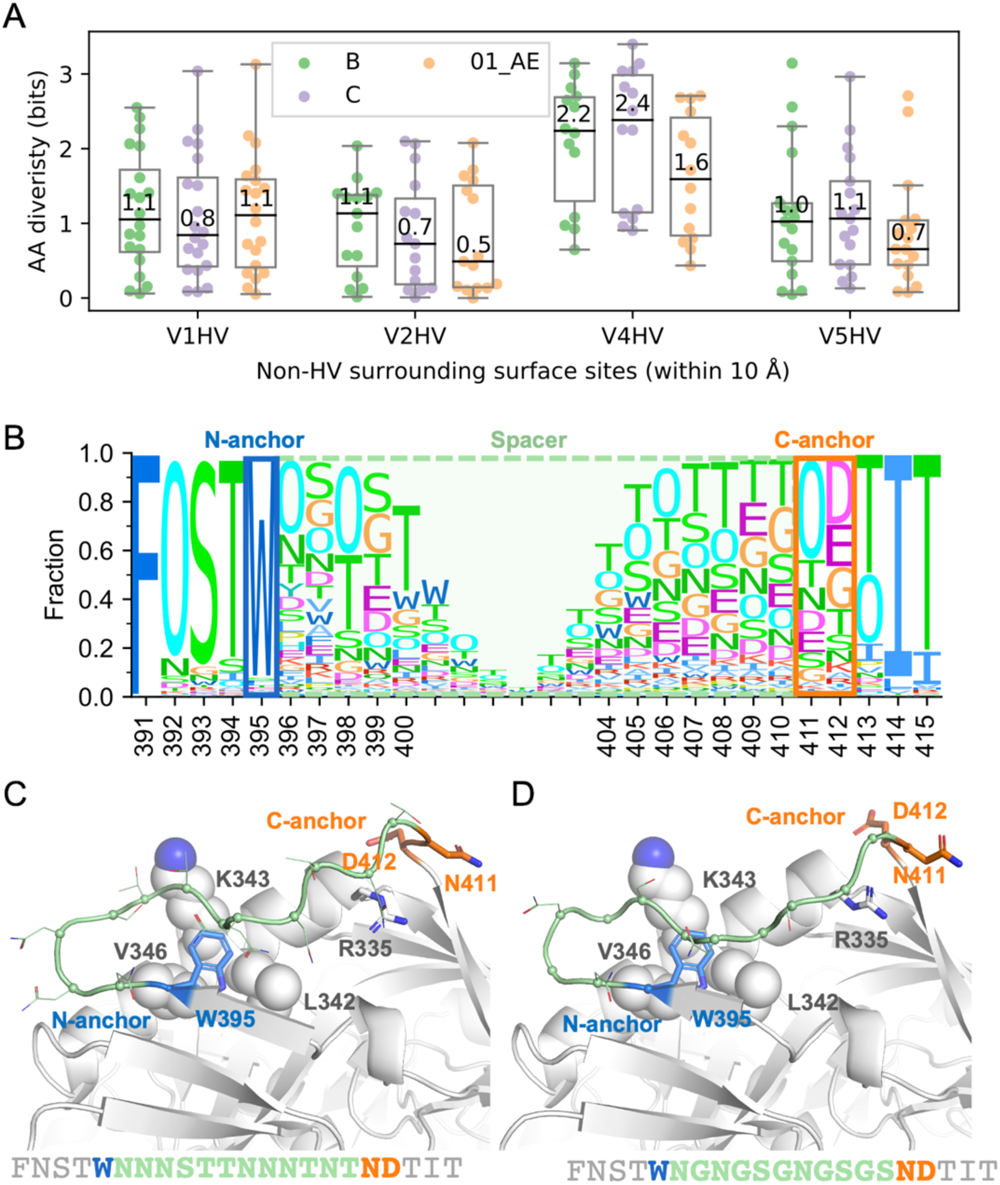
Redesigned V4 hypervariable (HV) loop of the subtype B Env consensus sequence. (A) The Shannon entropy of non-HV surrounding surface sites is shown for each loop and subtype/CRF, with median values reported. Key characteristics of the distribution, including 25th/75th percentile, are depicted by box plots. (B) Sequence logo of 2495 subtype B viruses around the V4HV loop, with the N-/C-anchor and spacer labeled. The height of each amino acid letter represents the frequency of that amino acid at a given position. The letter ‘O’ represents potential N-linked glycosylation sites. The consensus B Env with unmodified V4HV (C) and redesigned V4HV loop (D) modeled by AlphaFold2 are shown with the N-anchor, C-anchor and spacer sites colored blue, orange and light green, respectively. Non-HV residues that interact with anchor sites are shown as spheres or sticks.

### Redesigned HV loops improved accessibility to bnAb epitopes

To measure if the redesign improved exposure of bnAb epitopes, we calculated a depth index which corresponds to the closest distance between an epitope site and the antibody probe. The antibody probe is a sphere with a radius corresponding to an antibody’s Fv (=30 Å) that rolls around the Env structure (Fig. 5A). A short depth indicates that the site is more easily accessible to the antibody. We calculated the depth corresponding to the epitopes of two glycan supersite bnAbs, 10-1074 and PGT135, and two CD4bs bnAbs, VRC01 and 3BNC117, before and after the HV loop redesign. The epitope depth decreased for the four bnAb epitopes based on the structural model of the Env consensus of subtypes B, C, and CRF01 AE (Fig. 5B-D). The median depth decreased by 0.2 to 1.6 Å for glycan supersite antibody epitope sites, while the median depth decreased by 0.2 to 0.6 Å for CD4bs antibody epitope sites. Beside the depth, the clashes between HV loops and corresponding antibodies were also calculated to show the impact of the HV loop redesign on the antibody accessibility of bnAbs epitopes. The redesigned Env also alleviated the clashes between 10-1074 and PGT135 and V1HV, but with limited effect on the clashes between CD4bs antibodies and V5HV (Fig. 5E). The apex antibodies, PGT145 and VRC26.25, were not tested because their epitopes are quaternary and require modeling of the entire Env trimer, which is beyond our computational power.

**Fig 5.**
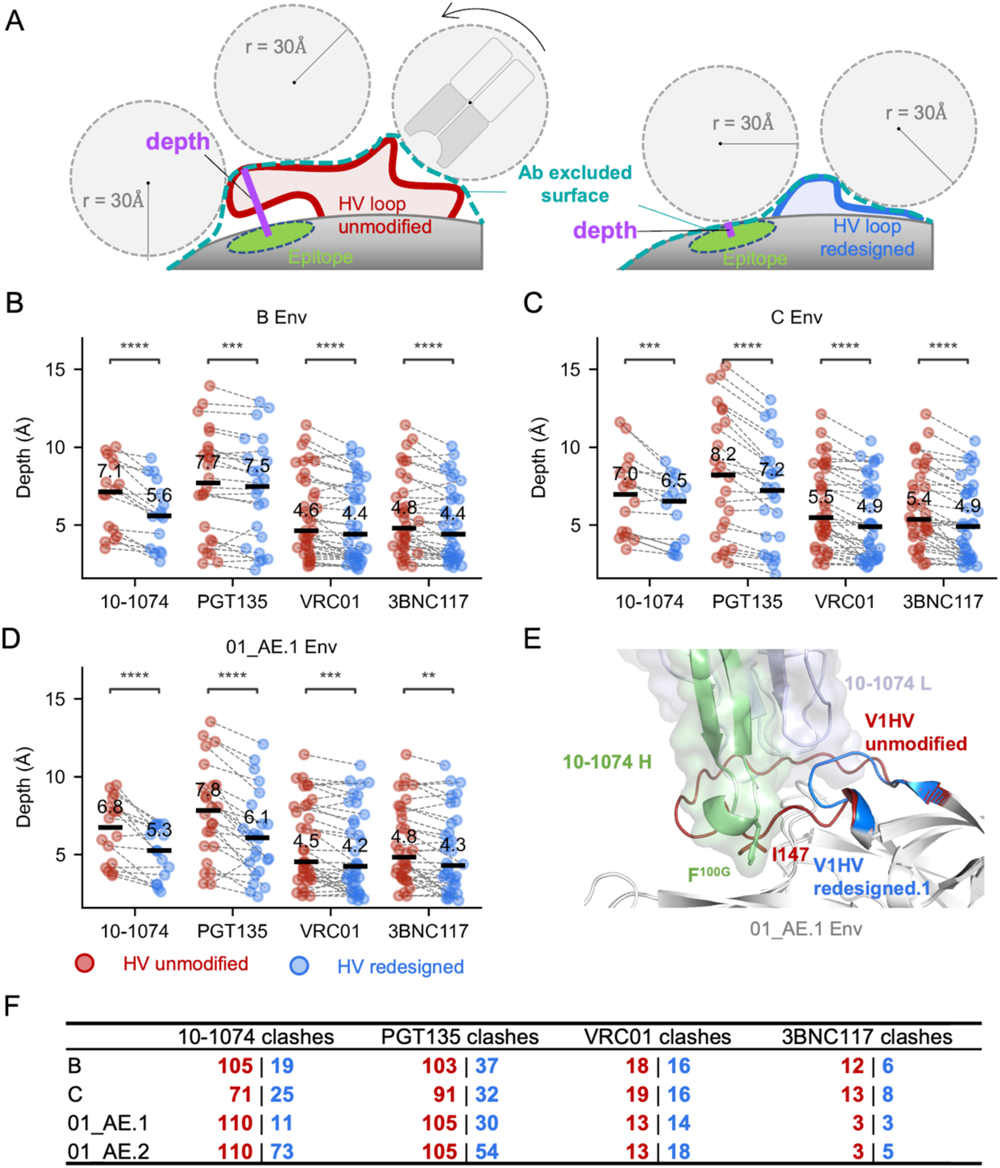
Impact of redesigned HV loops on the antibody accessibility of epitope sites. (A) Schematic showing how the minimum distance between an epitope site and the sphere probe (radius 30 Å) representing an antibody, defined as depth, is used to detect antibody accessibility differences of Env surface sites before (red) and after (blue) the HV loop redesign. (B)-(D) Comparison of the depth for four representative antibodies before and after the HV loop redesign. Median depth is labeled. The significance from paired, non-parametric Wilcoxon signed-rank tests is indicated as ‘**’, ‘***’, and ‘****’, for p ≤ 0.01, 0.001 and 0.0001, respectively. (E) Consensus CRF01_AE Env with unmodified and redesigned HV loops (AlphaFold2 prediction) are superimposed with Env in the 10-1074-Env complex. (F) Decrease in clashes between antibody atoms and Env for unmodified (red) and redesigned (blue) cognate HV loops.

We focused on the modeled structure of CRF01_AE V1HV because CRF01_AE Env have longer V1HV than those of subtypes B and C (Fig. 1B) and show high resistance to glycan supersite antibodies^47^. The consensus sequence had a ‘IGNITD’ motif (147-152) as its V1HV C-anchor sites (Fig. S3AB). AlphaFold2 predicted that I147 occupied the hydrophobic pocket formed by R327, K328, Y330 and P417, which corresponds to the site targeted by F^100G^ on the CDR-H3 of 10-1074 (Fig. 5E). This meant that the 10-1074 epitope was hidden with a median depth of 7.0 Å. After the HV loops redesign, the 24 AA-long V1HV loop (58.2 percentile in CRF01_AE) was replaced with a 9 AA-long loop (0.6 percentile in CRF01_AE, 01_AE.1) (Fig. S3C), hence, the median depth of 10-1074 epitope sites was reduced to 5.3 Å. The redesigned Env reduced the number of clashes with 10-1074 (Fig. 5E). A similar decrease in the number of clashes was observed for another glycan supersite bnAb, PGT135, but the impact was limited for CD4bs antibodies (VRC01 and 3BNC117) (Fig. 5F). The greater depth and clash reduction for glycan-supersite epitope sites near V1HV was likely due to the fact that V1HV loops were longer than other HV loops and the HV loop redesign removed more spacer residues than for other HV loops (Fig. 1A, Table 2). Because the ‘IGNITD’ motif is highly conserved in CRF01_AE viruses, another version of V1HV (01_AE.2) was designed to preserve the ‘IGNITD’ motif, which may be important for CRF01_AE specific bnAbs (Fig. S3D). This variant CRF01_AE.2 design also improved antibody accessibility for glycan super site antibodies albeit to a lesser extent, (Fig. 5F and S13).

### Improved antibody binding for Env with redesigned HV loops

To test if the HV loop redesign improves antibody binding to Env, we produced our redesigned consensus sequences with both unmodified and redesigned HV loops as gp140 glycoproteins. Both versions of the consensus subtype B and C, plus the unmodified CRF01_AE consensus and the redesigned V1HV with the conserved ‘IGNITD’ motif (01_AE.2) were tested for binding to plasma pools and monoclonal antibodies. The pooled plasma samples represented 13 cohorts of PLWH spanning diverse geographic areas and subtype/CRF distributions. Plasma antibodies across multiple clade-specific pools recognized the consensus B Env with redesigned HV loops at a slightly higher magnitude of binding compared to the Env with unmodified HV loops (Figure 6A). Similarly, monoclonal antibodies recognized the subtype B Env with redesigned loops at a higher magnitude of binding, especially for antibodies targeting the CD4bs, V2 apex and glycan supersite epitopes (Fig. 6B). Summary measures for binding of pooled plasma or monoclonal antibodies demonstrated minor but significantly higher recognition of the subtype B Env with redesigned HV loops when compared to the Env with unmodified loops (p≤0.001) (Fig. 6CD). Similar improvement of antibody binding was seen for the CRF01_AE consensus with redesigned HV loops (p≤0.00049), while the subtype C consensus proteins with or without redesigned loops showed no difference (Fig. 6CD, Fig. S5).

**Fig 6.**
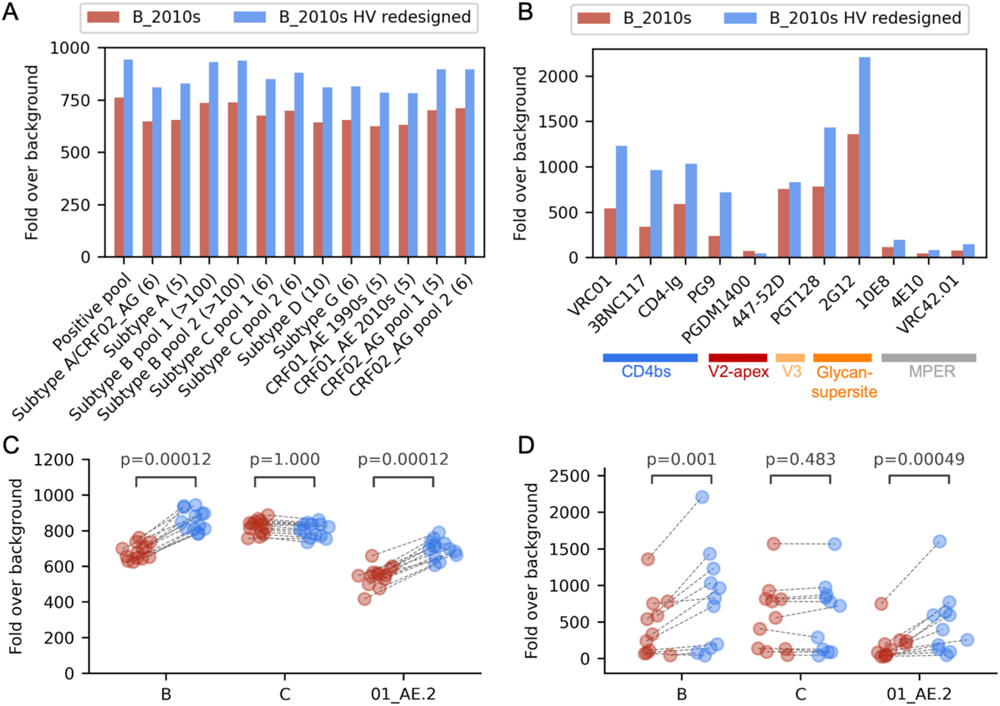
Improved antibody binding to subtypes B and C and CRF01_AE consensus Env with redesigned HV loops. Consensus Env sequences were expressed as gp140 glycoproteins. Comparison of the binding of (A) plasma pools from 13 cohorts of PLWH and (B) monoclonal antibodies to consensus B Env with unmodified (in red) and redesigned (in blue) HV loops. The number of plasma sample in each pool is shown in parenthesis in (A). The number of plasma samples in each pool is shown in parenthesis in panel A. Binding of plasma samples (C) and monoclonal antibodies (D) to the subtype B, C and CRF01_AE consensus Env with unmodified HV loops (red circles) and redesigned HV loops (blue circles) is compared using paired, non-parametric Wilcoxon signed-rank tests.

## Discussion

Here we redesigned the hypervariable loops V1HV, V2HV, V4HV and V5HV using a structural modeling strategy to optimize access to critical bnAb epitopes. The redesign was done for subtypes B and C and CRF01_AE. These subtypes/CRF were selected because large datasets of sequences were available (for the next most frequent subtype, A1, only 288 sequences were available). In addition, HIV-1 subtype B is the most thoroughly characterized subtype, subtype C is responsible for the majority of cases among PLWH and CRF01_AE is the virus that was circulating in the only HIV-1 vaccine trial where some efficacy was recorded^38^. We redesigned HV loops in Env consensus sequences that we had recently created; these consensus sequences reflected currently circulating viruses since they were derived only from sequences that were sampled after 2010. The LANL database recently updated HIV-1 consensus sequences using all sequences sampled since the beginning of the epidemic^48^, hence, these 2021 consensus sequences tend to be biased towards past sequences which are more numerous than sequences sampled in the past decade. We chose to restrict our consensus sequences to sequences sampled since 2010 to ensure an accurate representation of recently circulating viruses. We note that HIV-1 proteins that are available at the HIV Reagent Program (hivreagentprogram.org) and are commonly used as reagents in experimental assays were often produced two decades ago, Consensus sequences were chosen because they represent the theoretical most fit virus, and, as such, they are more likely than a natural Env sequence to present favorable features for structural integrity as well as ideal bnAb epitopes^32–35^. We showed that our updated contemporary consensus sequences were functional and that the consensus sequences with redesigned HV loops promoted antibody binding when compared to the unmodified updated consensus.

Our analysis of a large set of sequences showed that Env with long HV loops were more resistant to bnAbs targeting cognate epitopes than Env with shorter loops. This is consistent with previous findings showing that the combined length of V1HV and V2HV was correlated with virus sensitivity to glycan-supersite or V3-glycan bnAbs, and that the length of V5 was related to virus sensitivity to CD4bs antibodies^47^. Interestingly, our structural modeling uncovered a novel mechanism for CRF01_AE’s remarkable resistance to glycan-supersite antibodies such as 10-1074^47^. In addition to the absence of PNGS at site 332, I147 in CRF01_AE V1HV N-anchor sites was predicted to occupy the same location as the CDRH3 of 10-1074 (Fig. 5E and S1), a pocket composed of sites 328, 330, 415 and 417. In contrast, the pocket was not occupied by the modeled consensus B or C Env with unmodified V1HV. And PGT128, a V3-glycan antibody whose epitope does not occupy this pocket^13^ and is thus less occluded by the CRF01 AE V1HV, retains the capacity to neutralize half of the CRF01 AE viruses^47^. It is possible that other viruses with long V1HV use a similar mechanism to obstruct the binding of glycan-supersite bnAbs and divert antibody responses to these loops, highlighting that reducing the length of HV loops could play a key role in minimizing interference with bnAbs targeting adjacent epitopes. Importantly, the consensus sequences we used had HV loops of medium length due to how they are designed. Hence, redesigning the HV loop would likely have a more pronounced benefit in the context of a natural HIV-1 sequence.

Beside reducing HV loop length, we focused on reducing the immunogenicity of the HV loop length by using shorter spacers composed of less immunogenic amino acids. The redesigned spacers were composed of only glycine and serine residues except for the PNGS site. This is unlike natural HV loops, which consist of various AA. Our goal was to minimize strain-specific responses. Since glycine and serine lack antibody-recognizable side chains, the redesigned loops are less likely to act as decoys attracting strain-specific antibody responses. A recent study found that long V1V2s in CRF01_AE Env were associated with the more rapid development of autologous neutralization in participants from the RV217 cohort^49^; these results highlight the potential role of long HV loops in the development of strain specific responses – these strain-specific responses could be considered undesirable as autologous neutralization was not associated with the development of neutralization breadth in this cohort.

Importantly, we noted that HV loop length, particularly for V1, has not been an important consideration for vaccine inserts used in vaccine efficacy trials or under consideration for future trials (Table 1). Most of the sequences used in previous vaccines had HV loops that were longer than the median HV loop length in the population, suggesting that candidate inserts for future vaccine trials could be optimized to minimize the HV loop lengths. We want to emphasize that the HV loop redesign should be combined with other strategies. First, while bnAb epitopes near HV-loops were more accessible after our redesign, their accessibility may still be outcompeted by targeting of strain-specific glycan holes or of the base of the trimer, which is common when a soluble, stabilized prefusion Env is administered^50–54^. Thus, fixing strain-specific glycan holes, shielding the Env trimer base (e.g. by mounting it on a liposome^55^ or co-delivering it with Gag in an mRNA vaccine^45^) could also help improve the accessibility of bnAbs epitopes. Second, the epitope near the redesigned HV loops may also need to be optimized, though what constitutes an optimal epitope for the elicitation of bnAbs is debatable. We previously showed that bnAb epitopes were actually as diverse as non-bnAb epitopes among HIV-1 sequences; however, what distinguished the broadest bnAbs from the other nAbs targeting similar epitopes was their ability to focus on key epitope sites that were more conserved^6^. As a result, if rare mutations are found in key epitope sites, it would make sense to revert the mutated residues to the consensus AA at that site. Third, many unshielded epitopes are exposed on the post-fusion Env. Thus, for the HV loop modification to be effective, a stabilized prefusion Env trimer is required as the improved bnAbs accessibility may be insignificant in the post-fusion context. This can be achieved with the repair and stabilization procedure which systematically fixes rare mutations and stabilizes HIV-1 Env in the prefusion conformation^56,57^.

There are some limitations to our computational approach to assess antibody accessibility. First, the glycan shield is absent while we calculate the depth of the epitope and the clashes between Env and bnAbs. That might lead to an over estimation of the antibody accessibility both before and after the HV-loop redesign, as the occlusion from glycans is omitted. Second, we modeled one subunit in the prefusion Env trimer by AlphaFold2 and assumed the HV loop conformation from the top model. However, protein structure is dynamic, and long HV loops are frequently predicted with low confidence, as indicated by the pLDDT score. Third, when calculating the depth, a sphere with a radius of 30 Å was chosen as the antibody probe to simplify calculations, which is not necessarily a good approximation for Fv or Fab domain. As a result, our computational assessment should only be considered qualitatively correct in terms of capturing the antibody accessibility difference.

In summary, we redesigned new Env consensus sequences that reflect currently circulating sequences for subtypes B and C and CRF01_AE with modified HV loops. We demonstrated that long HV loops interfered with bnAbs targeting adjacent epitopes and that our modifications alleviated the interference of HV loops on bnAb epitopes, especially for the glycan-supersite bnAbs. Since half of all neutralizing antibodies target epitopes around the four optimized HV loops^22,23^, we propose that this design strategy could yield more effective antigens than those currently used as reagents or vaccine candidates. This optimization strategy can be integrated with other vaccine design techniques, such as the ‘repair and stabilize’ of Env in the prefusion conformation^56,57^, shielding the base of Env^45^, or the presentation of an optimum combination of Envs^58^, to create the next-generation of HIV-1 vaccines for eliciting broadly protective antibodies.

## Methods

### Dataset

Env AA sequences of subtypes B, C and CRF01_AE were retrieved from the Los Alamos National Laboratory HIV-1 sequence database on 2022-11-07. Sequences were retained if they had a complete open reading frame, information on the year and location of collection, and were not hypermutated or problematic (as defined by LANL). Non-independent sequences were removed so that the final dataset included one sequence per person. The IC50 of PGT145, VRC26.25, 10-1074, PGT135, VRC01 and 3BNC117 against viruses that had the corresponding Env sequences available were downloaded from CATNAP database as of 2022-10-04^46^. In total there were 549, 423, 782, 310, 1032 and 818 IC50s and associated Env sequences for PGT145, VRC26.25, 10-1074, PGT135, VRC01 and 3BNC117, respectively.

### HV loop analysis

The HXB2 genome annotation from LANL HIV Database (https://www.hiv.lanl.gov/content/sequence/HIV/MAP/annotation.html) was followed to define the start and end of HV loops, which were 132-152, 185-190, 396-410 and 460-467, for V1HV, V2HV, V4HV and V5HV, respectively. As an exception, V4HV was extended to include 395 and 411-412 anchor sites in the HV loop redesign. Structurally, the distance between the start and end of spacers was calculated as the distance between CA atoms before the N-terminal and after the C-terminal of the spacer loop. The HXB2 sequence was added and aligned to define the positions. MAFFT^59^ was used to align sequences. For the logo plot of HVs, HV loops were split from the middle: the first half was left-aligned to the start position while the second half was right-aligned to the end position.

### Structure prediction and analysis

We used local ColabFold^60^ which integrated AlphaFold-Multimer^61^ with other tools and databases^62–67^ to model unmodified and redesigned Env structures. During the prediction, one of the three subunits in the prefusion Env trimer is predicted by feeding one gp120 (31-511) and one gp41 (512-665) as input. The top model from five predictions was adopted for later analysis.

The non-HV surrounding surface sites of HV loops were defined as Env sites that were not part of the HV loop which have relative accessible surface area (RSA) greater or equal to 0.30 and have any atoms within 10 Å of any HV loop atoms in the predicted unmodified consensus B Env structure. The accessible surface area (ASA)^68^ was approximated by the Shrake-Rupley method^69^. In brief, evenly distributed points (n=256) were placed on spheres that were centered at atoms in the protein. The radius of the spheres was the sum of the atoms’ and the solvent probe’s radius (1.5 Å). The accessible surface was defined after filtering out points on a sphere that were within neighboring spheres. Then, RSA of a site was calculated as the ratio of ASA over the estimated maximum ASA^70^. The same approach was also used to estimate depth, only the probe radius was set to 30 Å, and the depth equals the minimal distance between a protein atom and the accessible surface minus the probe radius. Env structure 5FYJ^71^ was used for the ASA estimation. It was also used the illustration of HV loop and nearby bnAb epitopes in Figure 1.

The clashes between antibody and Env was determined as the number of antibody atoms within 2.5 Å of Env after superimposing the modeled Env on the Env in the solved Env-antibody complex. Bio.PDB^72^ in the Biopython^73^ was used to superimpose structures. The Env-antibody complexes used, listed as PDB code, were 5V8L^17^, 6VTT^18^, 6UDJ^74^, 4JM2^12^, 6V8X^75^ and 7PC2^76^ for PGT145, VRC26.25, 10-1074, PGT135, VRC01 and 3BNC117, respectively. PyMol^77^ was used for the structure visualization and figure generation. Hydrogen atoms, when present, were removed from the structure, in all analyses.

### Protein production and antibody binding

Uncleaved gp140 proteins were designed with R to S mutations in the gp120/gp41 furin cleavage site and incorporated the native leader peptide and full MPER sequence, followed by a short GGGS linker sequence and a C-terminal AviTag. Sequences, codon-optimized for expression in human cells, were synthesized (Genscript) and cloned into a custom pcDNA3.4 (ThermoFisher) expression vector. Proteins were expressed by transient transfections in Expi293F cells (ThermoFisher) according to the manufacturer’s instructions. Gp140 proteins were purified from clarified cell culture supernatants 4 days post-transfection using Galanthus nivalis lectin (GNL) affinity chromatography followed by a Q-sepharose (GE Healthcare) polishing step to remove host cell protein contaminants. All gp140s were purified to 90% purity or higher, as assessed by SDS-PAGE and Coomassie staining in reducing and non-reducing conditions. Antibodies VRC01 (PMID 20616231), 3BNC117 (PMID 27199429), 2G12 (PMID 7474069), PG9 (PMID 19729618), PGDM1400 (PMID 25422458), PGT128 (PMID21998254), 10E8 (PMID 23151583) and CD4-Ig (PMID 11805109) were produced in-house from public sequences as IgG1 by transfection in Expi293F cells and purified by protein A affinity chromatography. For antibody binding and characterization, gp140 proteins were coupled to uniquely coded carboxylated magnetic microspheres (Luminex Corp., Austin TX) per manufacturer’s protocol and as previously described and pooled into an antigen cocktail. Antibody binding and characterization was performed as previously described^78–80^ with minor modifications. The antigen cocktail was incubated with an array of heat-inactivated human plasma pool samples from various clade-specific cohorts of PLWH, or with monoclonal antibodies targeting various regions of HIV-1 Env. Samples were incubated for 2 hours at room temperature, washed then incubated for 1 hour with phycoerythrin labeled anti-human IgG Fc (Southern Biotech, Birmingham AL) detection antibody. After a final wash to remove unbound detection reagent, microspheres are resuspended in 40 μL sheath fluid (Luminex Corp) and data was collected on a Bio-Plex®3D Suspension Array system (Bio-Rad, Hercules CA) running xPONENT® v.4.2 (Luminex Corp). A multi-clade control plasma pool comprised of samples from HIV-1 Group O, subtypes B, C, and CRF01_AE, CRF02_AG and CRF03_AB (SeraCare, Milford MA) was used as a positive control (“Positive Pool”) and a pool of plasmas from people without HIV-1 (PWOH) served as a negative control. The figures correspond to monoclonal antibodies at a concentration of 10 ug/mL and pooled plasma at 1:100 dilution. Responses are reported as fold over PWOH for plasma samples and fold over the ZIKA virus specific monoclonal antibody MZ4^81^.

### QUANTIFICATION AND STATISTICAL ANALYSIS

Statistical analyses were performed using SciPy^82^. The non-paired, non-parametric Mann-Whitney U tests were used to compare the IC50 of short/long nearby or distant HV loops in Figure 2, S1 and S2. The paired, non-parametric Wilcoxon signed-rank tests were used to compare the depth difference before and after HV loop redesign (Figure 5) and the binding responses before and after the HV loop redesign with pooled plasma samples or bnAbs (Figure 6). Differences corresponding to p-values of less than 0.05 were considered as significant. Pandas^83^ and NumPy^84^ were used in the data analysis and Matplotlib^85^ was used to generate plots.

## Materials availability

Plasmids generated in this study will be available with a Material Transfer Agreement.

## Data availability

The data reported in this paper are available in the Supplemental material and at https://www.hivresearch.org/publication-supplements.

## Acknowledgments

We thank Johannes PM Langedijk and Lucy Rutten for helpful discussion, Michelle Zemil for technical assistance. This work was supported by a cooperative agreement between The Henry M. Jackson Foundation for the Advancement of Military Medicine, Inc., and the U.S. Department of the Army [WW81XWH-18-2-0040].

## Author contributions

H.B. and M.R. conceived the study, analyzed results and wrote the manuscript. H.B. designed and performed analyses. E.L. and Y.L. curated sequence alignments and designed consensus sequences. M.G.J. contributed to the characterization of redesigned Env. S.T, V.D., B.S., L.W., V.P., S.J.K., J.A.A. and S.V. produced Env proteins and mapped antibody binding responses.

## References

1. Mascola, J. R. & Montefiori, D. C. The role of antibodies in HIV vaccines. Annu. Rev. Immunol. 28, 413–444 (2010).

2. Saunders, K. O., Rudicell, R. S. & Nabel, G. J. The design and evaluation of HIV-1 vaccines. AIDS Lond. Engl. 26, 1293–1302 (2012).

3. Wibmer, C. K., Moore, P. L. & Morris, L. HIV broadly neutralizing antibody targets. Curr. Opin. HIV AIDS 10, 135–143 (2015).

4. Kwong, P. D. & Mascola, J. R. HIV-1 Vaccines Based on Antibody Identification, B Cell Ontogeny, and Epitope Structure. Immunity 48, 855–871 (2018).

5. Sok, D. & Burton, D. R. Recent progress in broadly neutralizing antibodies to HIV. Nat. Immunol. 19, 1179–1188 (2018).

6. Bai, H., Li, Y., Michael, N. L., Robb, M. L. & Rolland, M. The breadth of HIV-1 neutralizing antibodies depends on the conservation of key sites in their epitopes. PLoS Comput. Biol. 15, e1007056 (2019).

7. O’Connell, R. J., Kim, J. H. & Excler, J.-L. The HIV-1 gp120 V1V2 loop: structure, function and importance for vaccine development. Expert Rev. Vaccines 13, 1489–1500 (2014).

8. Moore, P. L., Gorman, J., Doria-Rose, N. A. & Morris, L. Ontogeny-based immunogens for the induction of V2-directed HIV broadly neutralizing antibodies. Immunol. Rev. 275, 217–229 (2017).

9. Kwong, P. D. et al. Structure of an HIV gp120 envelope glycoprotein in complex with the CD4 receptor and a neutralizing human antibody. Nature 393, 648–659 (1998).

10. Gristick, H. B. et al. Natively glycosylated HIV-1 Env structure reveals new mode for antibody recognition of the CD4-binding site. Nat. Struct. Mol. Biol. 23, 906–915 (2016).

11. Walker, L. M. et al. Broad neutralization coverage of HIV by multiple highly potent antibodies. Nature 477, 466–470 (2011).

12. Kong, L. et al. Supersite of immune vulnerability on the glycosylated face of HIV-1 envelope glycoprotein gp120. Nat. Struct. Mol. Biol. 20, 796–803 (2013).

13. Pejchal, R. et al. A potent and broad neutralizing antibody recognizes and penetrates the HIV glycan shield. Science 334, 1097–1103 (2011).

14. Buchacher, A. et al. Generation of human monoclonal antibodies against HIV-1 proteins; electrofusion and Epstein-Barr virus transformation for peripheral blood lymphocyte immortalization. AIDS Res. Hum. Retroviruses 10, 359–369 (1994).

15. Sok, D. et al. Recombinant HIV envelope trimer selects for quaternary-dependent antibodies targeting the trimer apex. Proc. Natl. Acad. Sci. U. S. A. 111, 17624–17629 (2014).

16. Liu, Q. et al. Quaternary contact in the initial interaction of CD4 with the HIV-1 envelope trimer. Nat. Struct. Mol. Biol. 24, 370–378 (2017).

17. Lee, J. H. et al. A Broadly Neutralizing Antibody Targets the Dynamic HIV Envelope Trimer Apex via a Long, Rigidified, and Anionic β-Hairpin Structure. Immunity 46, 690–702 (2017).

18. Gorman, J. et al. Structure of Super-Potent Antibody CAP256-VRC26.25 in Complex with HIV-1 Envelope Reveals a Combined Mode of Trimer-Apex Recognition. Cell Rep. 31, 107488 (2020).

19. Walker, L. M. et al. Broad and Potent Neutralizing Antibodies from an African Donor Reveal a New HIV-1 Vaccine Target. Science 326, 285–289 (2009).

20. Zhou, T. et al. Structural basis for broad and potent neutralization of HIV-1 by antibody VRC01. Science 329, 811–817 (2010).

21. Klein, F. et al. Somatic mutations of the immunoglobulin framework are generally required for broad and potent HIV-1 neutralization. Cell 153, 126–138 (2013).

22. Landais, E. et al. Broadly Neutralizing Antibody Responses in a Large Longitudinal Sub-Saharan HIV Primary Infection Cohort. PLoS Pathog. 12, e1005369 (2016).

23. Townsley, S. M. et al. B cell engagement with HIV-1 founder virus envelope predicts development of broadly neutralizing antibodies. Cell Host Microbe 29, 564–578.e9 (2021).

24. Srivastava, I. K., VanDorsten, K., Vojtech, L., Barnett, S. W. & Stamatatos, L. Changes in the immunogenic properties of soluble gp140 human immunodeficiency virus envelope constructs upon partial deletion of the second hypervariable region. J. Virol. 77, 2310–2320 (2003).

25. Silva de Castro, I., et al. Anti-V2 antibodies virus vulnerability revealed by envelope V1 deletion in HIV vaccine candidates. iScience 24, 102047 (2021).

26. Bontjer, I. et al. Comparative Immunogenicity of Evolved V1V2-Deleted HIV-1 Envelope Glycoprotein Trimers. PloS One 8, e67484 (2013).

27. Zolla-Pazner, S. et al. Rationally Designed Vaccines Targeting the V2 Region of HIV-1 gp120 Induce a Focused, Cross-Clade-Reactive, Biologically Functional Antibody Response. J. Virol. 90, 10993– 11006 (2016).

28. Jiang, X. et al. Rationally Designed Immunogens Targeting HIV-1 gp120 V1V2 Induce Distinct Conformation-Specific Antibody Responses in Rabbits. J. Virol. 90, 11007–11019 (2016).

29. Karch, C. P. et al. Design and characterization of a self-assembling protein nanoparticle displaying HIV-1 Env V1V2 loop in a native-like trimeric conformation as vaccine antigen. *Nanomedicine Nanotechnol*. Biol. Med. 16, 206–216 (2019).

30. Jumper, J. et al. Highly accurate protein structure prediction with AlphaFold. Nature 596, 583–589 (2021).

31. Kuiken, C., Korber, B. & Shafer, R. W. HIV sequence databases. *AIDS Rev.* 5, 52–61 (2003).

32. Korber, B. et al. Evolutionary and immunological implications of contemporary HIV-1 variation. Br. Med. Bull. 58, 19–42 (2001).

33. Gaschen, B. et al. Diversity considerations in HIV-1 vaccine selection. Science 296, 2354–2360 (2002).

34. Rolland, M. et al. Reconstruction and function of ancestral center-of-tree human immunodeficiency virus type 1 proteins. J. Virol. 81, 8507–8514 (2007).

35. Rolland, M. HIV-1 phylogenetics and vaccines. Curr. Opin. HIV AIDS 14, 227–232 (2019).

36. Gilbert, P. B. et al. Correlation between immunologic responses to a recombinant glycoprotein 120 vaccine and incidence of HIV-1 infection in a phase 3 HIV-1 preventive vaccine trial. J. Infect. Dis. 191, 666–677 (2005).

37. Pitisuttithum, P. et al. Randomized, double-blind, placebo-controlled efficacy trial of a bivalent recombinant glycoprotein 120 HIV-1 vaccine among injection drug users in Bangkok, Thailand. J. Infect. Dis. 194, 1661–1671 (2006).

38. Rerks-Ngarm, S. et al. Vaccination with ALVAC and AIDSVAX to prevent HIV-1 infection in Thailand. N. Engl. J. Med. 361, 2209–2220 (2009).

39. Barouch, D. H. et al. Mosaic HIV-1 vaccines expand the breadth and depth of cellular immune responses in rhesus monkeys. Nat. Med. 16, 319–323 (2010).

40. Hammer, S. M. et al. Efficacy trial of a DNA/rAd5 HIV-1 preventive vaccine. N. Engl. J. Med. 369, 2083–2092 (2013).

41. Bekker, L.-G. et al. Subtype C ALVAC-HIV and bivalent subtype C gp120/MF59 HIV-1 vaccine in low-risk, HIV-uninfected, South African adults: a phase 1/2 trial. *Lancet HIV* **5**, e366–e378 (2018).

42. Barouch, D. H. et al. Evaluation of a mosaic HIV-1 vaccine in a multicentre, randomised, double-blind, placebo-controlled, phase 1/2a clinical trial (APPROACH) and in rhesus monkeys (NHP 13-19). Lancet Lond. Engl. 392, 232–243 (2018).

43. LaBranche, C. C. et al. Neutralization-guided design of HIV-1 envelope trimers with high affinity for the unmutated common ancestor of CH235 lineage CD4bs broadly neutralizing antibodies. PLoS Pathog. 15, e1008026 (2019).

44. Gray, G. E. et al. Vaccine Efficacy of ALVAC-HIV and Bivalent Subtype C gp120-MF59 in Adults. N. Engl. J. Med. 384, 1089–1100 (2021).

45. Zhang, P. et al. A multiclade env-gag VLP mRNA vaccine elicits tier-2 HIV-1-neutralizing antibodies and reduces the risk of heterologous SHIV infection in macaques. Nat. Med. 27, 2234–2245 (2021).

46. Yoon, H. et al. CATNAP: a tool to compile, analyze and tally neutralizing antibody panels. Nucleic Acids Res. 43, W213–219 (2015).

47. Bricault, C. A. et al. HIV-1 Neutralizing Antibody Signatures and Application to Epitope-Targeted Vaccine Design. Cell Host Microbe 25, 59–72.e8 (2019).

48. Linchangco, G. V., Foley, B. & Leitner, T. Updated HIV-1 Consensus Sequences Change but Stay Within Similar Distance From Worldwide Samples. Front. Microbiol. 12, 828765 (2021).

49. Kuriakose Gift, S., et al. Evolution of Antibody Responses in HIV-1 CRF01_AE Acute Infection: Founder Envelope V1V2 Impacts the Timing and Magnitude of Autologous Neutralizing Antibodies. J. Virol. e0163522 (2023) doi:10.1128/jvi.01635-22.

50. Klasse, P. J. et al. Sequential and Simultaneous Immunization of Rabbits with HIV-1 Envelope Glycoprotein SOSIP.664 Trimers from Clades A, B and C. PLoS Pathog. **12**, e1005864 (2016).

51. Nogal, B. et al. Mapping Polyclonal Antibody Responses in Non-human Primates Vaccinated with HIV Env Trimer Subunit Vaccines. Cell Rep. 30, 3755–3765.e7 (2020).

52. Reiss, E. I. M. M. et al. Fine-mapping the immunodominant antibody epitopes on consensus sequence-based HIV-1 envelope trimer vaccine candidates. NPJ Vaccines 7, 152 (2022).

53. Hu, J. K. et al. Murine Antibody Responses to Cleaved Soluble HIV-1 Envelope Trimers Are Highly Restricted in Specificity. J. Virol. 89, 10383–10398 (2015).

54. McCoy, L. E. et al. Holes in the Glycan Shield of the Native HIV Envelope Are a Target of Trimer-Elicited Neutralizing Antibodies. Cell Rep. 16, 2327–2338 (2016).

55. Shao, S. et al. Functionalization of cobalt porphyrin-phospholipid bilayers with his-tagged ligands and antigens. Nat. Chem. 7, 438–446 (2015).

56. Rutten, L. et al. A Universal Approach to Optimize the Folding and Stability of Prefusion-Closed HIV-1 Envelope Trimers. Cell Rep. 23, 584–595 (2018).

57. Rawi, R. et al. Automated Design by Structure-Based Stabilization and Consensus Repair to Achieve Prefusion-Closed Envelope Trimers in a Wide Variety of HIV Strains. Cell Rep. 33, 108432 (2020).

58. Lewitus, E., Hoang, J., Li, Y., Bai, H. & Rolland, M. Optimal sequence-based design for multi-antigen HIV-1 vaccines using minimally distant antigens. PLoS Comput. Biol. 18, e1010624 (2022).

59. Katoh, K. & Standley, D. M. MAFFT multiple sequence alignment software version 7: improvements in performance and usability. Mol. Biol. Evol. 30, 772–780 (2013).

60. Mirdita, M. et al. ColabFold: Making Protein folding accessible to all. Nat. Methods (2022) doi:10.1038/s41592-022-01488-1.

61. Evans, R., et al. Protein complex prediction with AlphaFold-Multimer. *bioRxiv* (2021) doi:10.1101/2021.10.04.463034v1.

62. Mirdita, M., Steinegger, M. & S“oding, J. MMseqs2 desktop and local web server app for fast, interactive sequence searches. Bioinformatics 35, 2856–2858 (2019).

63. Steinegger, M. et al. HH-suite3 for fast remote homology detection and deep protein annotation. BMC Bioinform 20, 473 (2019).

64. Eastman, P. et al. OpenMM 7: Rapid development of high performance algorithms for molecular dynamics. PLOS Comput Biol 13, (2017).

65. Berman, H., Henrick, K. & Nakamura, H. Announcing the worldwide Protein Data Bank. Nat. Struct. Biol. vol. 10 980 (2003).

66. Mirdita, M. et al. Uniclust databases of clustered and deeply annotated protein sequences and alignments. Nucleic Acids Res 45, D170–D176 (2017).

67. Mitchell, A. L. et al. MGnify: the microbiome analysis resource in 2020. Nucleic Acids Res (2019) doi:10.1093/nar/gkz1035.

68. Lee, B. & Richards, F. M. The interpretation of protein structures: estimation of static accessibility. J. Mol. Biol. 55, 379–400 (1971).

69. Shrake, A. & Rupley, J. A. Environment and exposure to solvent of protein atoms. Lysozyme and insulin. J. Mol. Biol. 79, 351–371 (1973).

70. Rost, B. & Sander, C. Conservation and prediction of solvent accessibility in protein families. Proteins 20, 216–226 (1994).

71. Stewart-Jones, G. B. E. et al. Trimeric HIV-1-Env Structures Define Glycan Shields from Clades A, B, and G. Cell 165, 813–826 (2016).

72. Hamelryck, T. & Manderick, B. PDB file parser and structure class implemented in Python. Bioinforma. Oxf. Engl. 19, 2308–2310 (2003).

73. Cock, P. J. A. et al. Biopython: freely available Python tools for computational molecular biology and bioinformatics. Bioinforma. Oxf. Engl. 25, 1422–1423 (2009).

74. Schommers, P. et al. Restriction of HIV-1 Escape by a Highly Broad and Potent Neutralizing Antibody. Cell 180, 471–489.e22 (2020).

75. Henderson, R. et al. Disruption of the HIV-1 Envelope allosteric network blocks CD4-induced rearrangements. Nat. Commun. 11, 520 (2020).

76. Lorin, V. et al. Epitope convergence of broadly HIV-1 neutralizing IgA and IgG antibody lineages in a viremic controller. J. Exp. Med. 219, e20212045 (2022).

77. Schrödinger, LLC. The PyMOL Molecular Graphics System, Version 1.8. (2015).

78. Tomaras, G. D. et al. Initial B-cell responses to transmitted human immunodeficiency virus type 1: virion-binding immunoglobulin M (IgM) and IgG antibodies followed by plasma anti-gp41 antibodies with ineffective control of initial viremia. J. Virol. 82, 12449–12463 (2008).

79. Brown, E. P. et al. High-throughput, multiplexed IgG subclassing of antigen-specific antibodies from clinical samples. J. Immunol. Methods 386, 117–123 (2012).

80. Mdluli, T. et al. RV144 HIV-1 vaccination impacts post-infection antibody responses. PLoS Pathog. 16, e1009101 (2020).

81. Dussupt, V. et al. Potent Zika and dengue cross-neutralizing antibodies induced by Zika vaccination in a dengue-experienced donor. Nat. Med. 26, 228–235 (2020).

82. Virtanen, P. et al. SciPy 1.0: fundamental algorithms for scientific computing in Python. Nat. Methods 17, 261–272 (2020).

83. McKinney, W. Data Structures for Statistical Computing in Python. in *Proceedings of the 9th Python in Science Conference* (eds. Walt, S. van der & Millman, J.) 56–61 (2010). doi:10.25080/Majora-92bf1922-00a.

84. Harris, C. R. et al. Array programming with NumPy. Nature 585, 357–362 (2020).

85. Hunter, J. D. Matplotlib: A 2D Graphics Environment. Comput. Sci. Eng. 9, 90–95 (2007).

