## Supplemental Information for "Contemporary HIV-1 consensus Env with redesigned hypervariable loops promote antibody binding"

### Supplementary material

#### Table S1. Summary measures for V1, V2, V4 and V5 HV loop lengths.

| | V1HV | | | V2HV | | | V4HV | | | V5HV | | |
|---|---|---|---|---|---|---|---|---|---|---|---|---|
| | B | C | 01_AE | B | C | 01_AE | B | C | 01_AE | B | C | 01_AE |
| count | 2495 | 1503 | 849 | 2495 | 1503 | 849 | 2495 | 1503 | 849 | 2495 | 1503 | 849 |
| mean | 22.8 | 20.0 | 23.8 | 8.0 | 9.1 | 8.5 | 12.5 | 9.2 | 8.6 | 9.2 | 9.2 | 8.5 |
| std | 6.0 | 6.1 | 4.1 | 3.7 | 3.5 | 3.6 | 3.7 | 4.4 | 3.5 | 2.1 | 2.4 | 1.8 |
| min | 3 | 5 | 7 | 0 | 0 | 0 | 0 | 0 | 0 | 0 | 1 | 2 |
| 25% | 19 | 16 | 22 | 6 | 6 | 6 | 11 | 6 | 7 | 8 | 8 | 7 |
| 50% | 22 | 20 | 23 | 7 | 9 | 8 | 12 | 10 | 9 | 9 | 9 | 8 |
| 75% | 26 | 24 | 26 | 10 | 11 | 10 | 15 | 12 | 11 | 10 | 10 | 9 |
| max | 44 | 40 | 40 | 26 | 23 | 27 | 27 | 24 | 22 | 19 | 21 | 17 |

#### Table S2. Summary measures for the number of PNGS in V1, V2, V4 and V5 HV loops.

| | V1HV | | | V2HV | | | V4HV | | | V5HV | | |
|---|---|---|---|---|---|---|---|---|---|---|---|---|
| | B | C | 01_AE | B | C | 01_AE | B | C | 01_AE | B | C | 01_AE |
| count | 2495 | 1503 | 849 | 2495 | 1503 | 849 | 2495 | 1503 | 849 | 2495 | 1503 | 849 |
| mean | 2.6 | 2.6 | 3.1 | 1.2 | 1.2 | 1.1 | 2.4 | 2.0 | 1.8 | 1.5 | 1.3 | 1.6 |
| std | 1.0 | 1.0 | 1.0 | 0.6 | 0.6 | 0.6 | 0.9 | 1.0 | 0.7 | 0.6 | 0.6 | 0.5 |
| min | 0 | 0 | 0 | 0 | 0 | 0 | 0 | 0 | 0 | 0 | 0 | 0 |
| 25% | 2 | 2 | 3 | 1 | 1 | 1 | 2 | 1 | 1 | 1 | 1 | 1 |
| 50% | 3 | 2 | 3 | 1 | 1 | 1 | 2 | 2 | 2 | 2 | 1 | 2 |
| 75% | 3 | 3 | 4 | 2 | 2 | 1 | 3 | 3 | 2 | 2 | 2 | 2 |
| max | 7 | 7 | 7 | 5 | 4 | 5 | 5 | 5 | 5 | 4 | 4 | 3 |

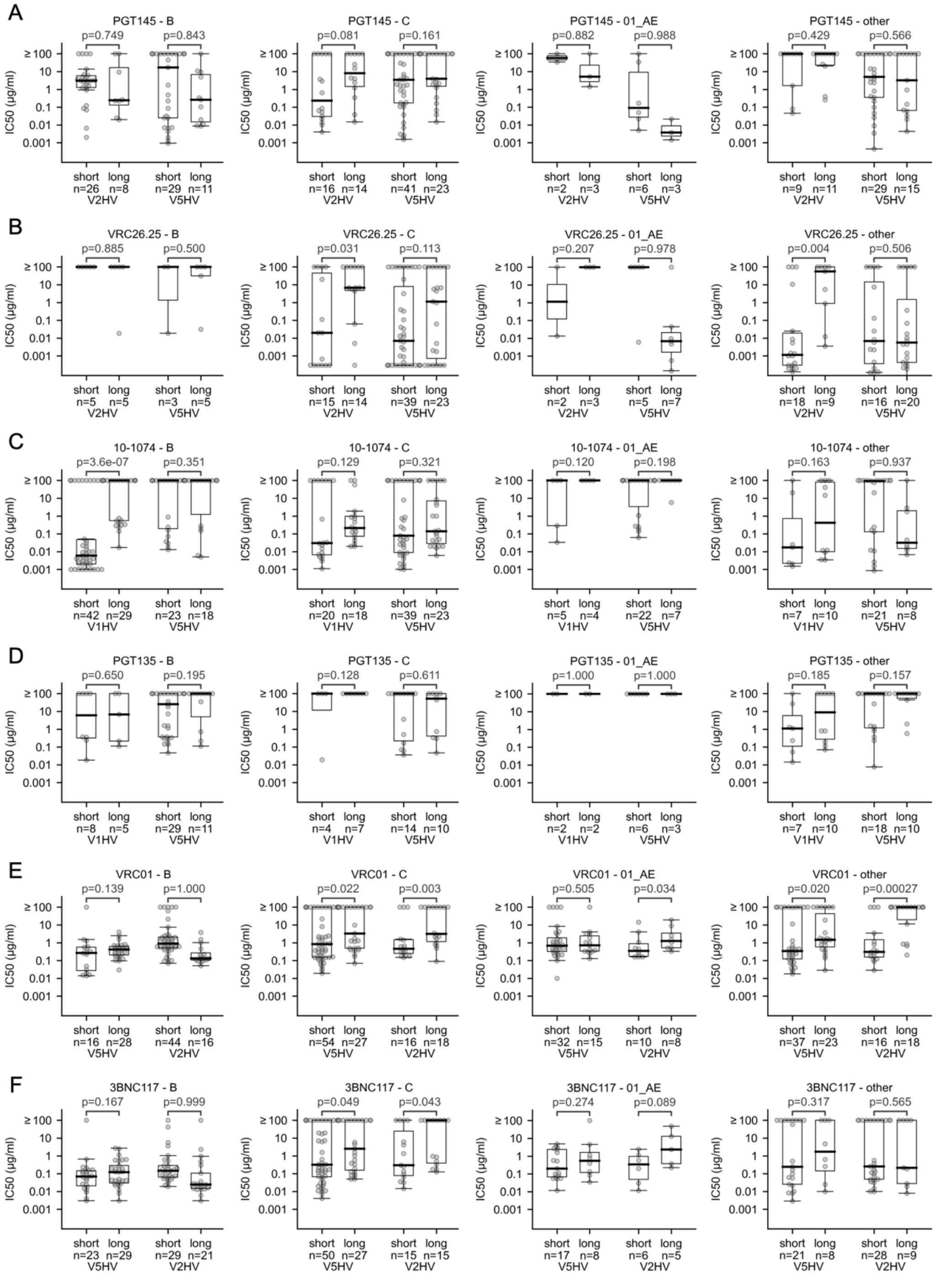


**Fig S1. Comparison of the bnAb sensitivity for HIV-1 Env with short (≤ 5 percentile) or long (≥ 95 percentile) hypervariable (HV) loops of different subtypes.** Comparisons were performed on two V2 apex-targeting antibodies whose epitope are close to V2HV loops: PGT145 (A) and VRC26.25 (B); two glycan supersite antibodies whose epitopes are close to V1HV loops: 10-1074 (C) and the PGT135 (D); and two CD4 binding site antibodies with epitopes close to the V5HV loops: VRC01 (E) and 3BNC117 (F). For each panel, comparisons of short and long HV loops are shown for the HV loop adjacent to (left) and remote from (right) the Ab epitope. The p-values correspond to non-paired, non-parametric Mann-Whitney U tests.

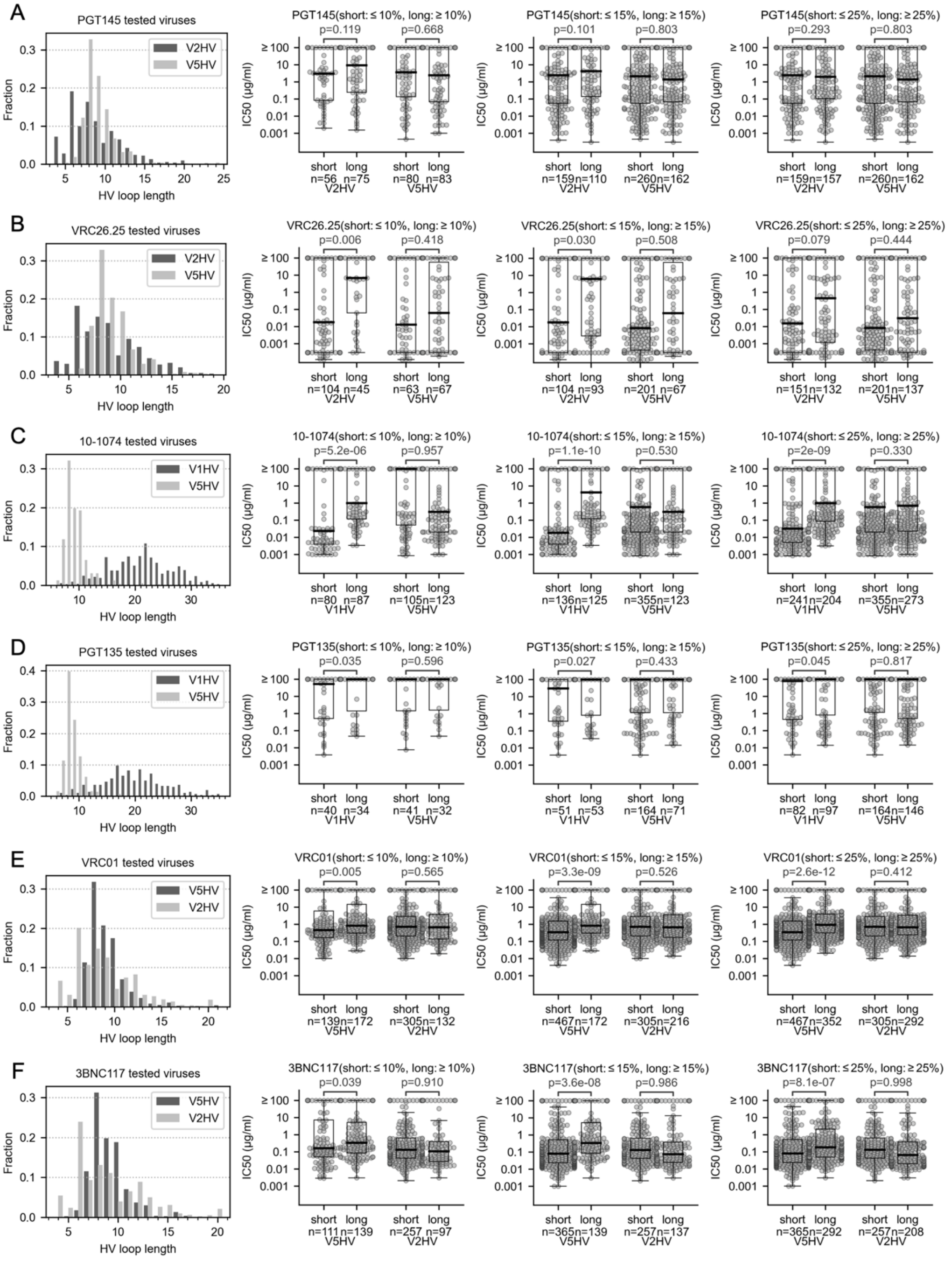


**Fig S2. Comparison of the bnAb sensitivity for HIV-1 Env with different cutoff from shortest or longest hypervariable (HV) loops.** The comparisons are based on six antibodies: PGT145 (A), VRC26.25 (B), 10-1074 (C), PGT135 (D), VRC01 (E) and 3BNC117 (F). The first column shows the distribution of HV loop length for tested viruses in CATNAP. The second to fourth columns are the comparison of sensitivity between viruses with short/long adjacent and distant HV loops, with cutoff of 10%,15% or 25%, respectively. The number of viruses can be the same for different cutoffs, as viruses with the cutoff length can correspond to 40% of all the tested Env sequences (e.g. V5HV of PGT135 tested viruses). The p-values correspond to non-paired, non-parametric Mann-Whitney U test.

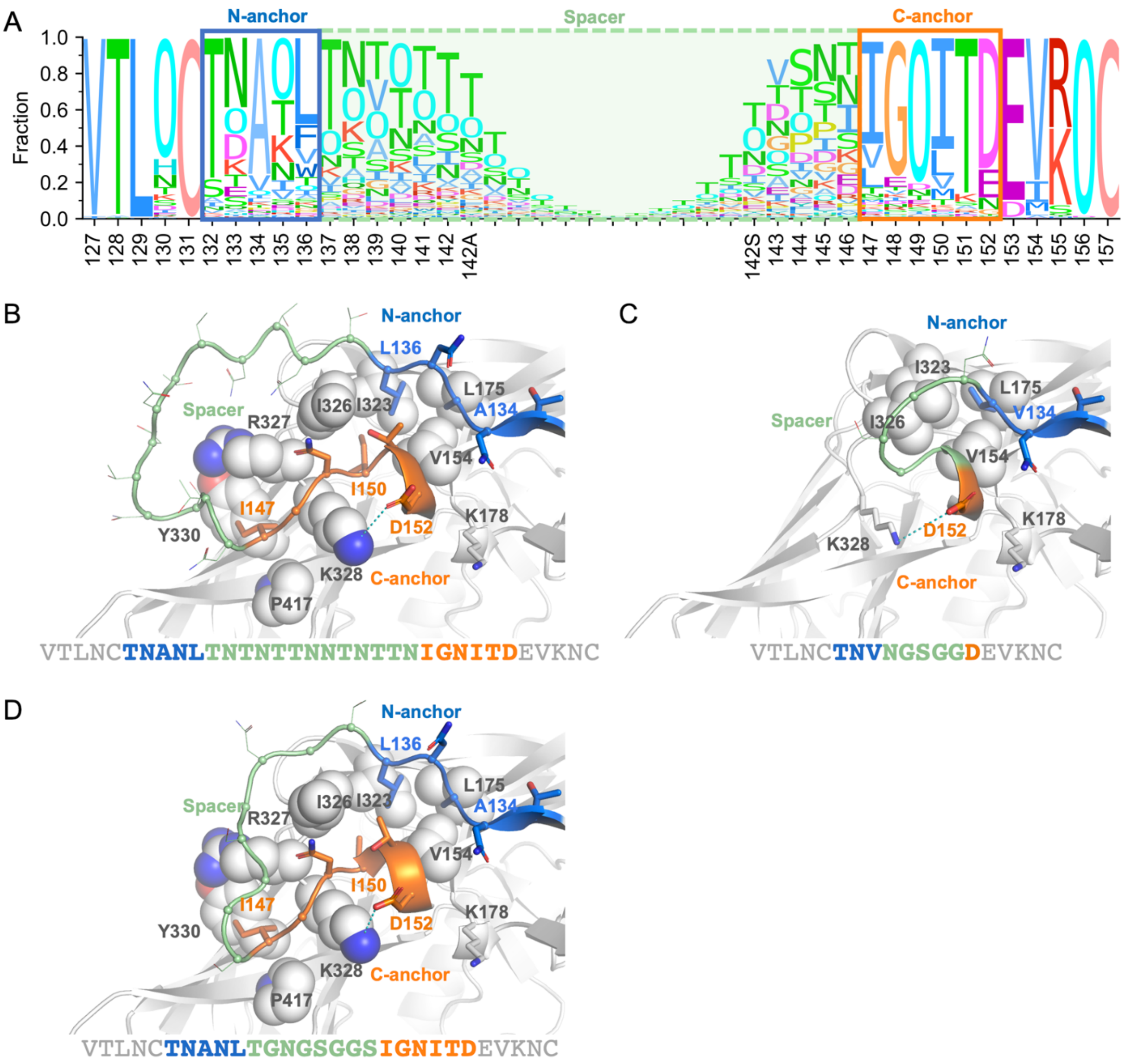


**Fig S3. Redesigned V1 hypervariable loop of the CRF01_AE consensus sequence.** (A) Sequence logo of 849 CRF01_AE sequences around the V1HV loop, with the N-/C-anchors and spacer indicated by boxes. The letter ‘O’ indicates a potential N-linked glycosylation site. The consensus CRF01_AE Env with unmodified V1HV loop (B) and redesigned V1HV loop (C, D) modeled by AlphaFold2 are shown with the N-anchor, C-anchor and spacer sites colored blue, orange and light green, respectively. ‘IGNITD’, the conserved C-anchor sites are replaced by ‘D’ to improve antibody accessibility to the 10-1074 epitope (C) or kept as is in the expectation of better recognition by CRF01_AE antibodies (D). Non-HV residues that interact with anchor sites are shown as spheres or sticks.

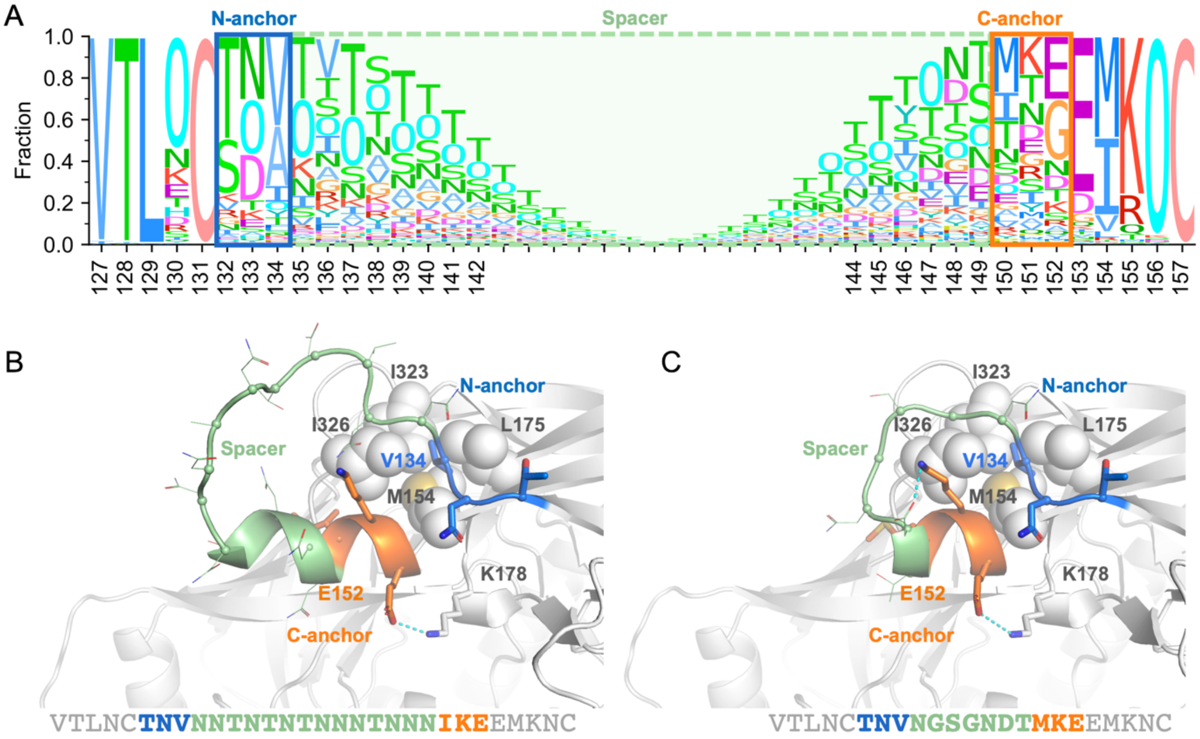


**Fig S4. Redesigned V1 hypervariable loop of the subtype C consensus sequence.** (A) Sequence logo of 1503 subtype C sequences around the V1HV loop, with the N-/C-anchors and spacer indicated by boxes. The letter ‘O’ indicates a potential N-linked glycosylation site. The consensus subtype C Env with unmodified V1HV loop (B) and redesigned V1HV loop (C) modeled by AlphaFold2 are shown with the N-anchor, C-anchor and spacer sites colored blue, orange and light green, respectively. Non-HV residues that interact with anchor sites are shown as spheres or sticks.

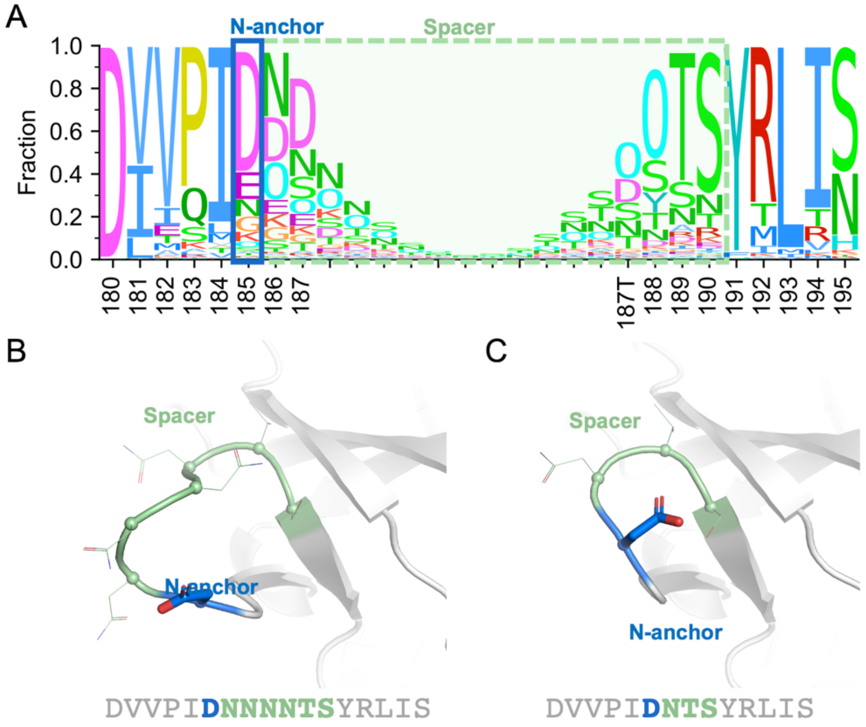


**Fig S5. Redesigned V2 hypervariable loop of the subtype B consensus sequence.** (A) Sequence logo of 2495 subtype B viruses around the V2HV loop, with the N-anchor and spacer indicated by boxes. The letter 'O' indicates a potential N-linked glycosylation site. The consensus subtype B Env with unmodified V2HV loop (B) and redesigned V2HV loop (C) modeled by AlphaFold2 are shown with the N-anchor and spacer sites colored blue and light green, respectively.

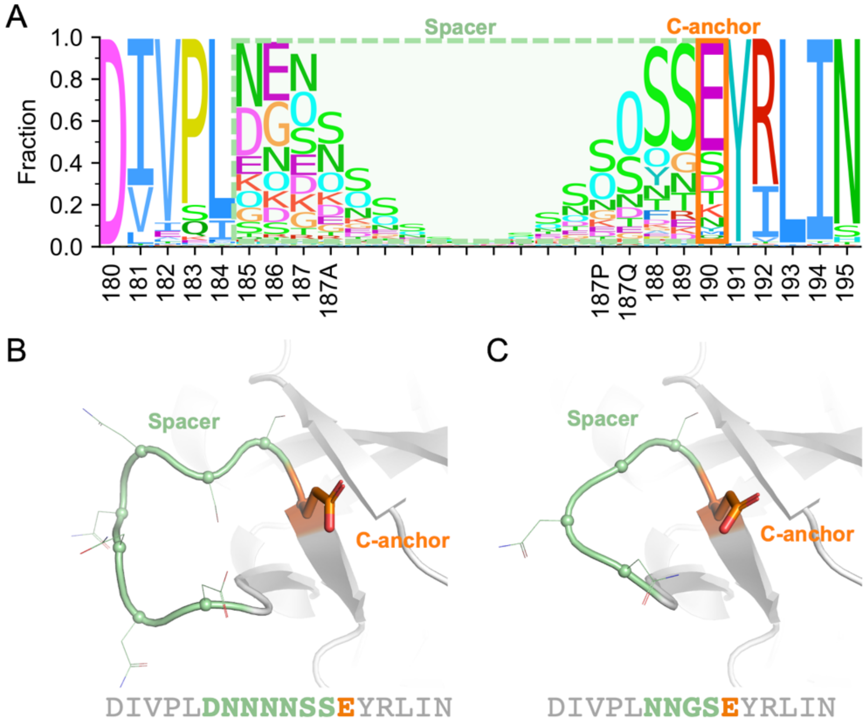


**Fig S6. Redesigned V2 hypervariable loop of the subtype C consensus sequence.** (A) Sequence logo of 1503 subtype C sequences around the V2HV loop, with the C-anchors and spacer indicated by boxes. The letter 'O' indicates a potential N-linked glycosylation site. The consensus subtype C with unmodified V2HV loop (B) and redesigned V2HV loop (C) modeled by AlphaFold2 are shown with the C-anchor and spacer sites colored orange and light green, respectively.

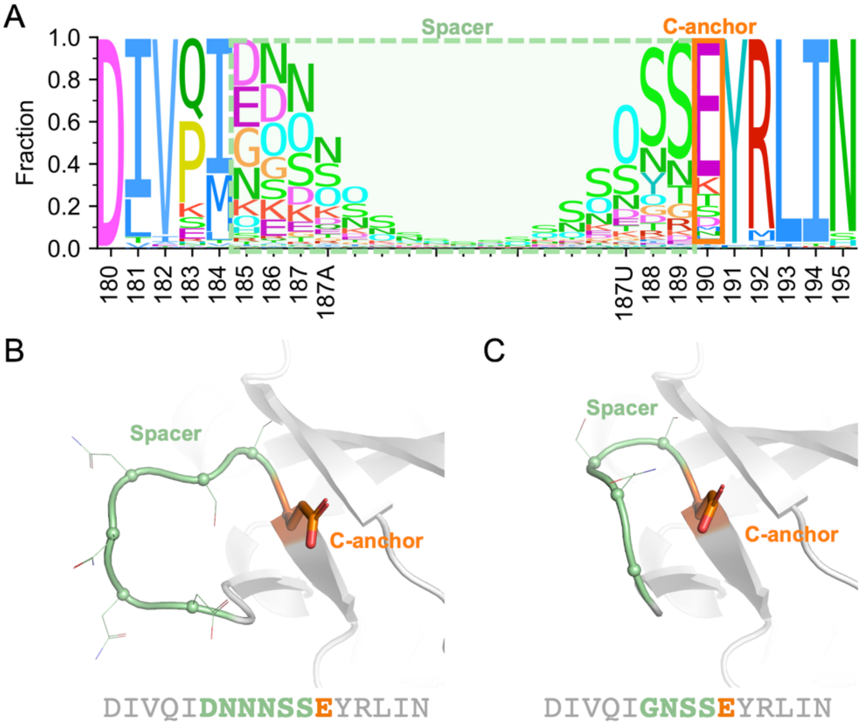


**Fig S7. Redesigned V2 hypervariable loop of CRF01_AE consensus sequence.** (A) Sequence logo of 849 CRF01_AE sequences around the V2HV loop, with the C-anchors and spacer indicated by boxes. The letter ‘O’ indicates a potential N-linked glycosylation site. The consensus CRF01_AE Env with unmodified V2HV loop (B) and redesigned V2HV loop (C) modeled by AlphaFold2 are shown with the C-anchor and spacer sites colored orange and light green, respectively.

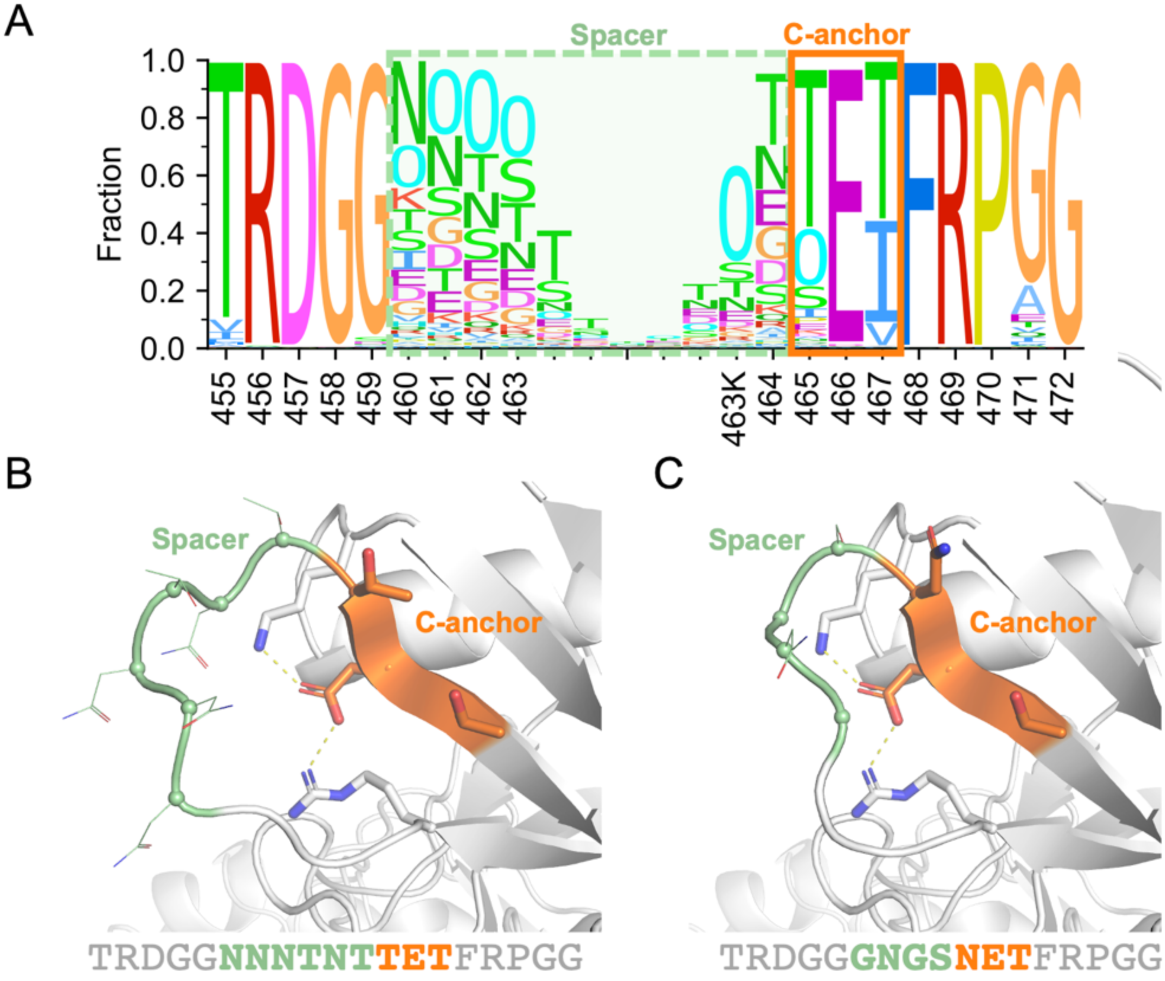


**Fig S8. Redesigned V5 hypervariable loop of the subtype B consensus sequence.** (A) Sequence logo of 2495 subtype B sequences around the V5HV loop, with the C-anchor and spacer indicated by boxes. The letter 'O' indicates a potential N-linked glycosylation site. The consensus subtype B Env with unmodified V5HV loop (B) and redesigned V5HV loop (C) modeled by AlphaFold2 are shown with the C-anchor and spacer sites colored orange and light green, respectively. Non-HV residues that interact with anchor sites are shown as sticks.

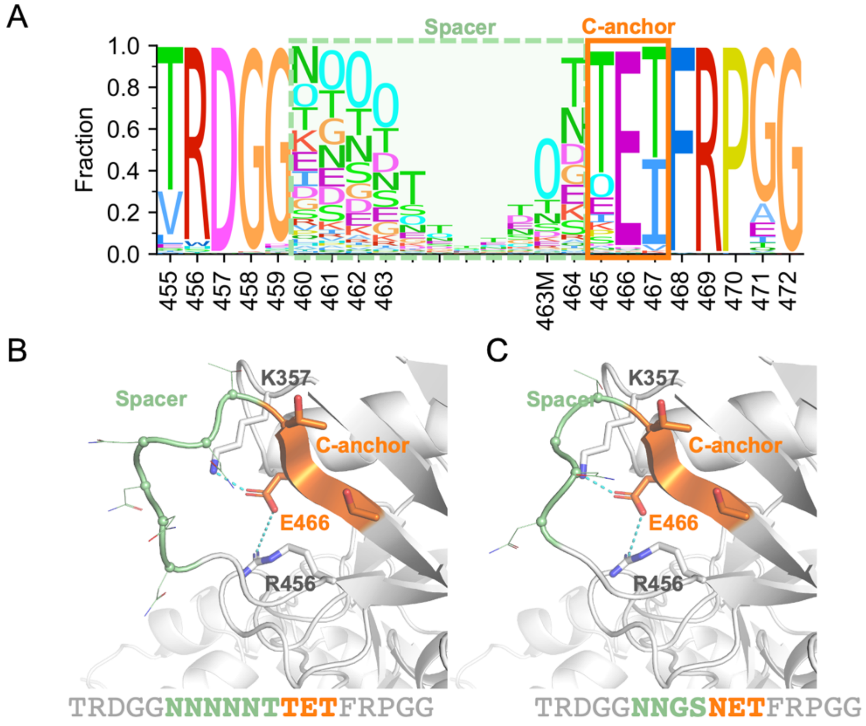


**Fig S9. Redesigned V5 hypervariable loop of the subtype C consensus sequence.** (A) Sequence logo of 1503 subtype C sequences around the V5HV loop, with the C-anchor and spacer indicated by boxes. The letter 'O' indicates a potential N-linked glycosylation site. The consensus subtype C Env with unmodified V5HV loop (B) and redesigned V5HV loop (C) modeled by AlphaFold2 are shown with the C-anchor and spacer sites colored orange and light green, respectively. Non-HV residues that interact with anchor sites are shown as sticks.

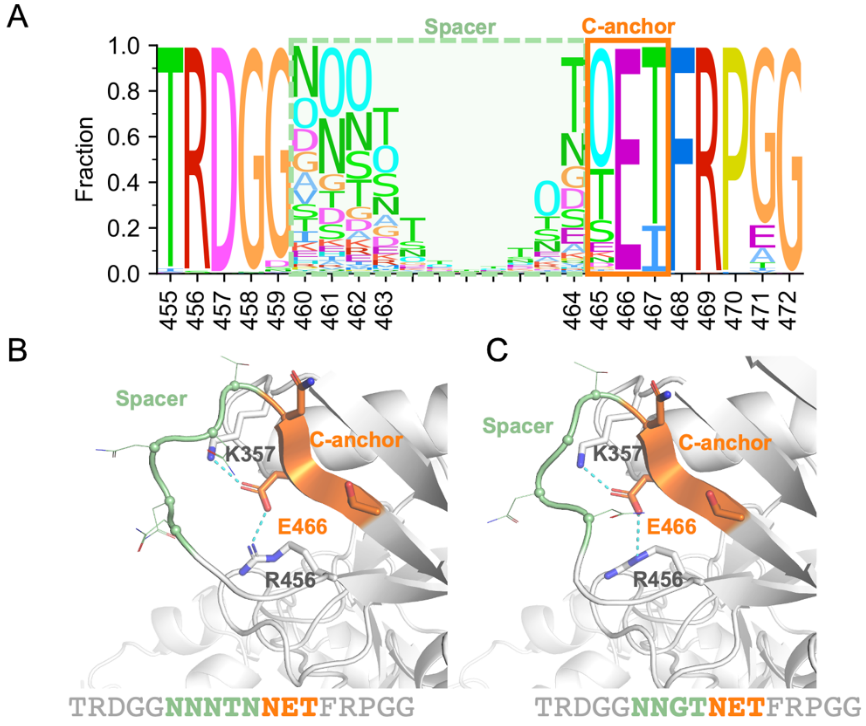


**Fig S10. Redesigned V5 hypervariable loop of CRF01_AE consensus sequence.** (A) Sequence logo of 849 CRF01_AE sequences around the V5HV loop, with the C-anchor and spacer indicated by boxes. The letter 'O' indicates a potential N-linked glycosylation site. The consensus CRF01_AE Env with unmodified V5HV loop (B) and redesigned V5HV loop (C) modeled by AlphaFold2 are shown with the C-anchor and spacer sites colored orange and light green, respectively. Non-HV residues that interact with anchor sites are shown as sticks.

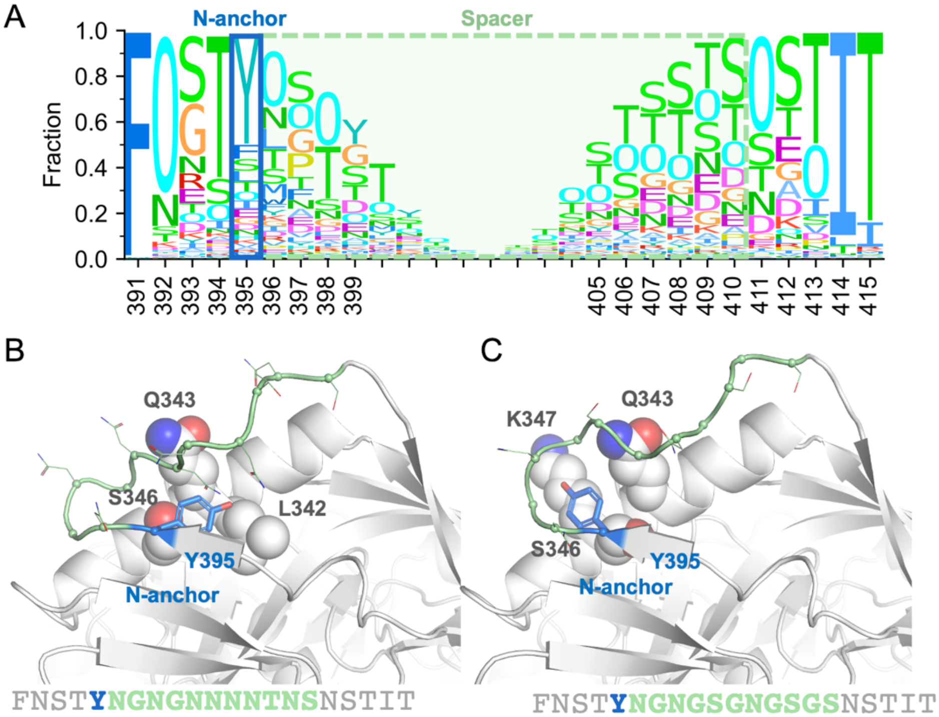


**Fig S11. Redesigned V4 hypervariable loop of the subtype C consensus sequence.** (A) Sequence logo of 1503 subtype C sequences around the V4HV loop, with the N-anchors and spacer indicated by boxes. The letter 'O' indicates a potential N-linked glycosylation site. The consensus subtype C Env with unmodified V4HV loop (B) and redesigned V4HV loop (C) modeled by AlphaFold2 are shown with the N-anchor and spacer sites colored blue and light green, respectively. Non-HV residues that interact with anchor sites are shown as spheres.

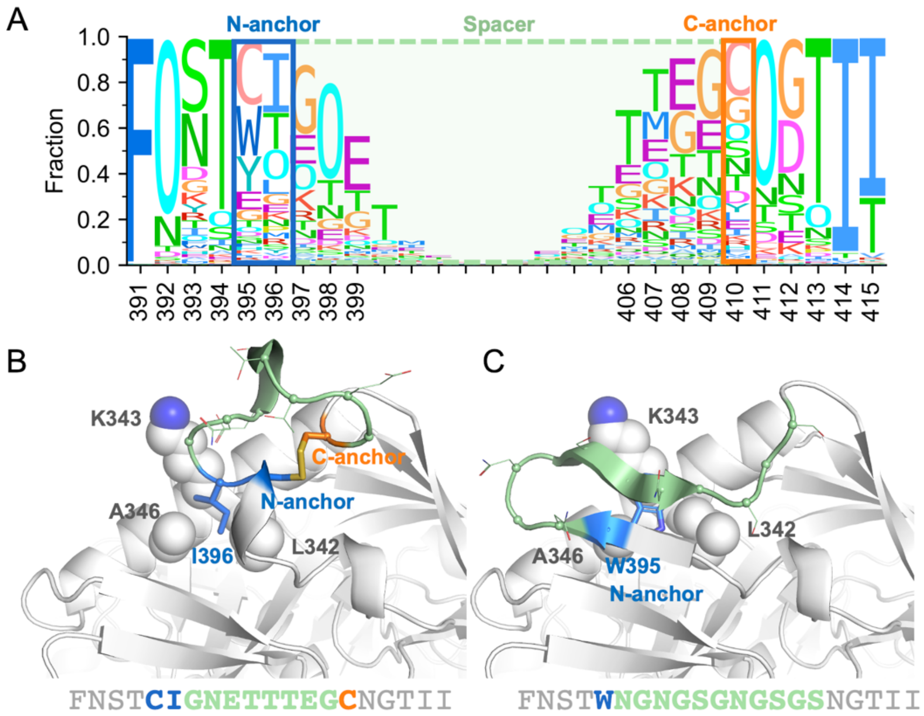


**Fig S12. Redesigned V4 hypervariable loop of CRF01_AE consensus sequence.** (A) Sequence logo of 849 CRF01_AE sequences around the V4HV loop, with the N-/C-anchors and spacer indicated by boxes. The letter 'O' indicates a potential N-linked glycosylation site. The consensus CRF01_AE Env with unmodified V4HV loop (B) and redesigned V4HV loop (C) modeled by AlphaFold2 are shown with the N-anchor, C-anchor and spacer sites colored blue, orange and light green, respectively. Non-HV residues that interact with anchor sites are shown as spheres.

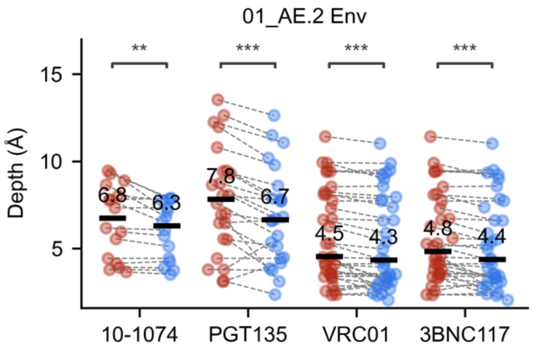


**Fig S13. The depth for four representative antibodies before and after the HV loop redesign shows an improvement in antibody accessibility with redesigned loops (01_AE.2).** The definition of depth is illustrated in Figure 5. The significance from Wilcoxon signed-rank tests is indicated as '**' and '***', for p ≤ 0.01 and 0.001, respectively.

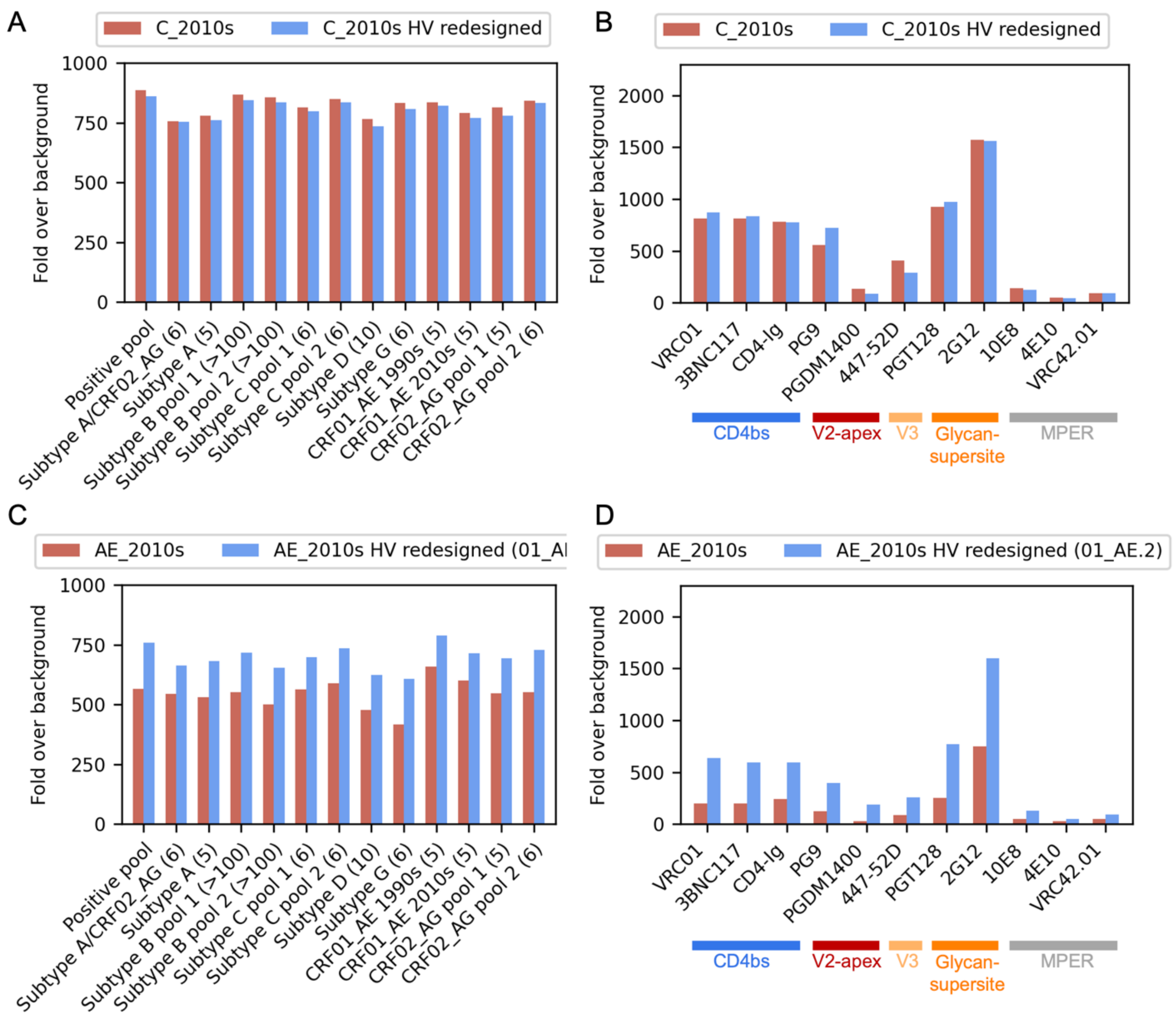


**Fig S14. Antibody binding to redesigned consensus Env from subtype C and CRF01_AE.** Consensus Env sequences with unmodified and redesigned HV loops were expressed as gp140 proteins to compare binding to 13 plasma pools combined from different cohorts of PWOH (A, C) and to monoclonal antibodies (B, D). The number of plasma samples in each pool is shown in parenthesis in panel A. The consensus C and CRF01_AE Env with redesigned HV loops are colored in blue and Env with unmodified HV loops are colored in red.
